# Heart cockle shells transmit sunlight to photosymbiotic algae using bundled fiber optic cables and condensing lenses

**DOI:** 10.1101/2022.10.28.514291

**Authors:** Dakota E. McCoy, Dale H. Burns, Elissa Klopfer, Liam K. Herndon, Babatunde Ogunlade, Jennifer A. Dionne, Sönke Johnsen

## Abstract

Many animals convergently evolved photosynthetic symbioses. In bivalves, giant clams (Cardiidae: Tridacninae) gape open to irradiate their symbionts, but heart cockles (Cardiidae: Fraginae) stay closed because sunlight passes through transparent windows in their shells. Here, we show that heart cockles (*Corculum cardissa* and spp.) use biophotonic adaptations to transmit sunlight for photosynthesis. Heart cockles transmit 11-62% of photosynthetically active radiation (mean=31%) but only 5-28% of potentially harmful UV radiation (mean=14%) to their symbionts. Beneath each window, microlenses condense light to penetrate more deeply into the symbiont-rich tissue. Within each window, aragonite forms narrow fibrous prisms perpendicular to the surface. These bundled “fiber optic cables’’ project images through the shell with a resolution of >100 lines/mm. Parameter sweeps show that the aragonite fibers’ size (∼1µm diameter), morphology (long fibers rather than plates), and orientation (along the optical c-axis) transmit more light than many other possible designs. Heart cockle shell windows are thus: (i) the first instance of fiber optic cable bundles in an organism to our knowledge; (ii) a second evolution, with epidermal cells in angiosperm plants, of condensing lenses for photosynthesis; and (iii) a photonic system that efficiently transmits useful light while protecting photosymbionts from UV radiation.

## Introduction

Photosynthesis is the engine that powers much of life on our planet. Many researchers focus on photosynthesis in plants, algae, and cyanobacteria. However, animals also harness sunlight indirectly through symbiotic partnerships ^1,2^. These animals– including certain reef-building corals, sponges, and bivalves--rely on a suite of optical adaptations to provide light for their photosymbionts ^2^. These little-studied optical adaptations can reveal how hosts and symbionts coevolve, how photosynthetic systems arose convergently, and how we could design bio-inspired technologies.

Photosymbiotic bivalves are a particularly interesting group despite receiving less attention than corals. Bivalvia is a class of primarily shelled molluscs that includes clams, oysters, mussels, and other marine and freshwater clades. Bivalves appear to have evolved obligate photosymbiosis at least twice, in two subfamilies within the family Cardiidae: the giant clams (Tridacninae) and the heart cockles (Fraginae ^3–6^; see ^7^ for review of other opportunistic symbioses in bivalves). To survive, giant clams and heart cockles require the products of photosynthesis from their dinoflagellate partners; correspondingly, to photosynthesize, the dinoflagellates require sunlight.

Bivalves in these two groups, having hard and normally opaque shells, had to evolve techniques to let light irradiate their soft tissues and the photosynthetic algae within. Giant clams solve this problem by gaping their shell open to bathe their mantle in downwelling sunlight. They further refined their photosynthetic capabilities through layered iridocytes that forward-scatter photosynthetically active radiation, back-reflect other wavelengths, and absorb ultraviolet (UV) light that is then re-emitted at longer wavelengths through fluorescence ^8–10^. Certain cardiid bivalves (e.g. *Clioniocardium nuttalli*, Clinocardiinae, and *Fragum* spp., Fragiinae) also temporarily gape their shells to expose mantle tissue, and, in some cases, extend mantle tissue out of and over the shell surface ^7,11^. But there is an alternative solution that does not require that a clam expose its soft mantle to predation, UV radiation, and other dangers: windows in the shell.

Heart cockles (*Corculum cardissa* and spp.) evolved transparent windows in their otherwise opaque shell to allow light to reach their symbionts ^12–16^. The common name “heart cockle” typically refers to *Corculum cardissa*, but the genus *Corculum* contains seven recognized species– although *C. cardissa* is the only well-studied species from a biophotonics perspective ^14,17,18^. Here, we used the general term heart cockle to refer to the *Corculum* spp. shells studied herein.

Heart cockles have shells with asymmetric valves (Figure 1), and are found partially buried in sand or corals, at water depths of 0.5-10 meters ^3,13^. Sunlight irradiates the photosynthetic dinoflagellates *Symbiodinium corculorum* (Symbiodiniaceae) within the mantle, gills, and foot ^3,12,13^. If placed in the shade, a heart cockle will move to the sun. If the sun-facing half of a heart cockle is covered with sand or mud, it will use its foot to sweep its shell clean ^16^.

**Figure 1:**
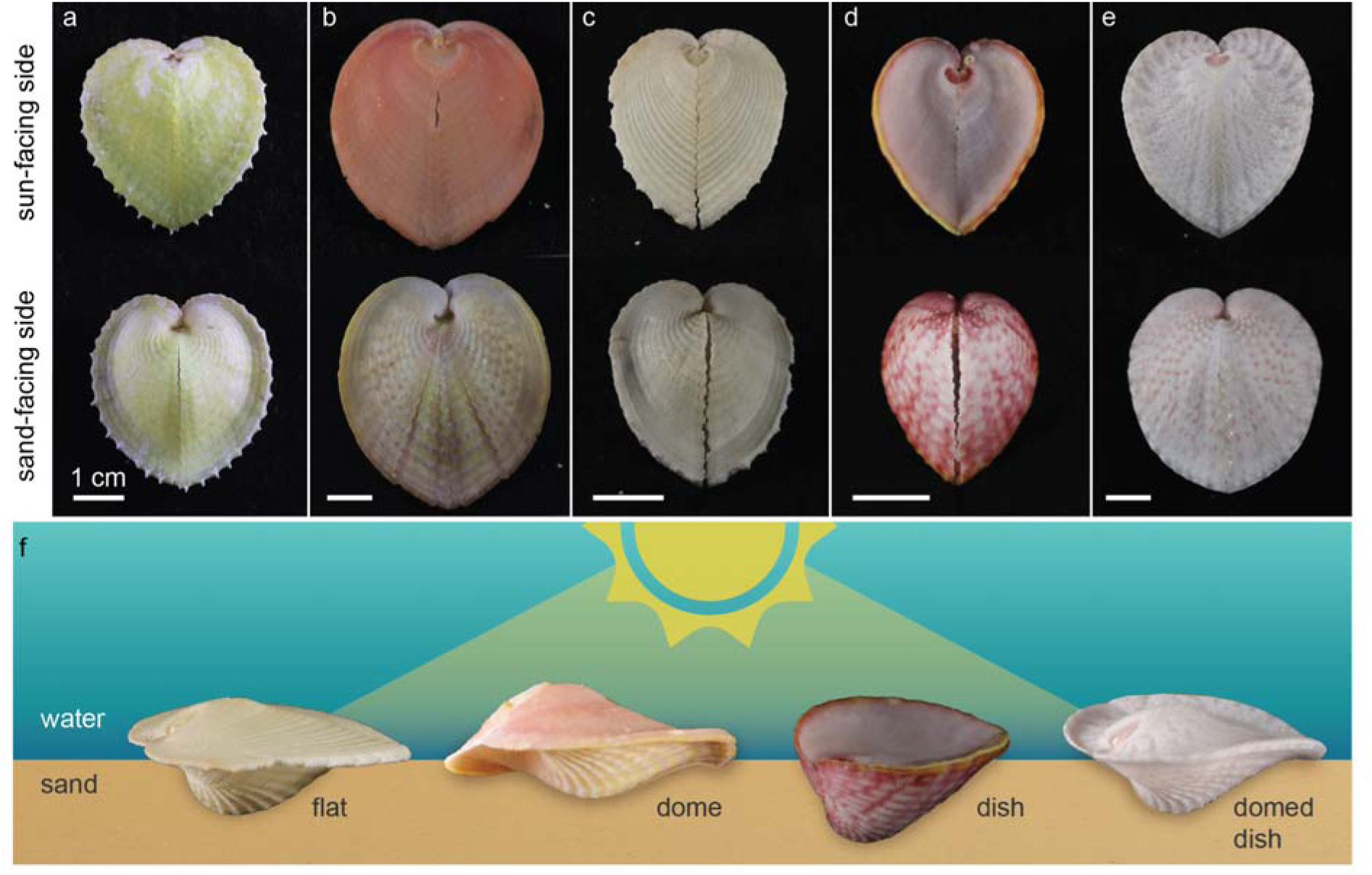
Heart cockles (*Corculum cardissa* and *Corculum* spp.) are asymmetrical, photosymbiotic bivalves. **(a-e)** Heart cockles come in many sizes, shapes, and colors across **(a-c)** *Corculum cardissa* and the seven other recognized species of *Corculum*, e.g., **(d)** *Corculum roseum* and **(e)** *Corculum lorenzi*. **(f)** The sun-facing side of heart cockle shells varies from a flattened, expanded pancake shake (“flat”; *Corculum cardissa*) to an arched dome (“dome”; *Corculum cardissa*) to a cup-shaped dish (“dish”; *Corculum roseum*) and combinations of these shapes (e.g., “domed dish”; *Corculum lorenzi*). Illustration in **(f)** is credited to Nuria Melisa Morales Garcia.

The shells of heart cockles are made from aragonite ^19^, a crystalline form of calcium carbonate (CaCO_3_) widely used by molluscs and cnidarians to make hard parts, and an organic matrix that directs the growth of the crystals ^20^. Two features distinguish the windows from the opaque regions of the shell. First, the aragonite within the windows forms elongated, fibrous crystals ^13–16,21^, the function of which has not yet been resolved ^22^. In opaque regions of the shell, the aragonite tends to be planar and crossed in orientation (termed “crossed lamellar” ^15,16^). Second, the windows are contiguous with transparent bumps on the inner surface of the shell that have been proposed to function as dispersing ^23,24^ or condensing ^15^ lenses ^13–16,21^.

Here, we show that heart cockles use natural nanophotonics in shell windows that screen UV radiation but transmit ample sunlight for photosynthesis. We apply a suite of photonic experiments and simulations to characterize these shell windows. The windows transmit more than twice as much photosynthetically useful sunlight as they do harmful UV radiation (mean=31% vs 14%; range=11-62% vs 5-28%). In shell windows, aragonite forms bundled fiber optic cables that project high resolution images ( >100 lines/mm); beneath the windows, aragonite condensing lenses focus light. Parameter sweeps show that the morphology and orientation of the aragonite fiber optic cables sit at a rough evolutionary optimum compared to other possibilities. To our knowledge, heart cockles are the only known evolution of bundled fiber optics and are a second evolution of condensing lenses for photosynthesis (alongside angiosperm plants). These bivalves illustrate the physics of photosynthesis across the tree of life and may inspire new engineered nanotechnologies.

## Results and Discussion

### Heart cockles have transparent windows that transmit photosynthetically-active radiation and screen out UV radiation

Heart cockles (*Corculum cardissa* and *Corculum* spp.) are asymmetrical bivalves whose shells vary in shape and color (Figure 1). Some individual shells are strongly yellow (Figure 1a), orange-pink (Figure 1b), or pink (Figure 1d), while others are white with only hints of color (Figure 1c,e). Heart cockles are antero-posteriorly flattened and form a heart-shaped outline when viewed from above (Figure 1a-e). The sun-facing side of heart cockle shells differs in profile from the sand-facing side, an asymmetry characteristic to heart cockles (Figure 1f). The sun-facing side is flat in some shells while other shells are shaped like a dome, dish, or combination of dome and dish (Figure 1f).

Transparent windows are arranged radially on the sun-facing half of the shell, with the exact shape and arrangement of the windows varying from triangular (Figure 2a) to radial stripes (Figure 2b) or even mosaics (Figure 2c-d). Approximately 50% of each shell surface is windowed (estimated by pixel brightness in imageJ). When viewed from above, triangular shell windows average 0.71 mm^2^ in area (n=50 windows, Figure 2a) and stripes averaged 0.58 mm in width (n=14, Figure 2b). Using a lapidary saw, we cut nearly-flat 1 cm^2^ fragments from heart cockle shell specimens (i.e., a section of shell with minimal natural curvature). Then, we suspended these fragments in seawater in a cuvette to measure transmission through the sun-facing side and the more opaque sand-facing side. The sun-facing side fragments included many tiny, evenly-spaced windows (Figure 2), so the 1 cm^2^ fragments are a good representation of the overall shell. We also measured transmission through individual shell windows polished to a width of 300μm (Supplementary Figure 1), the width used in our simulations (Figure 6).

**Figure 2:**
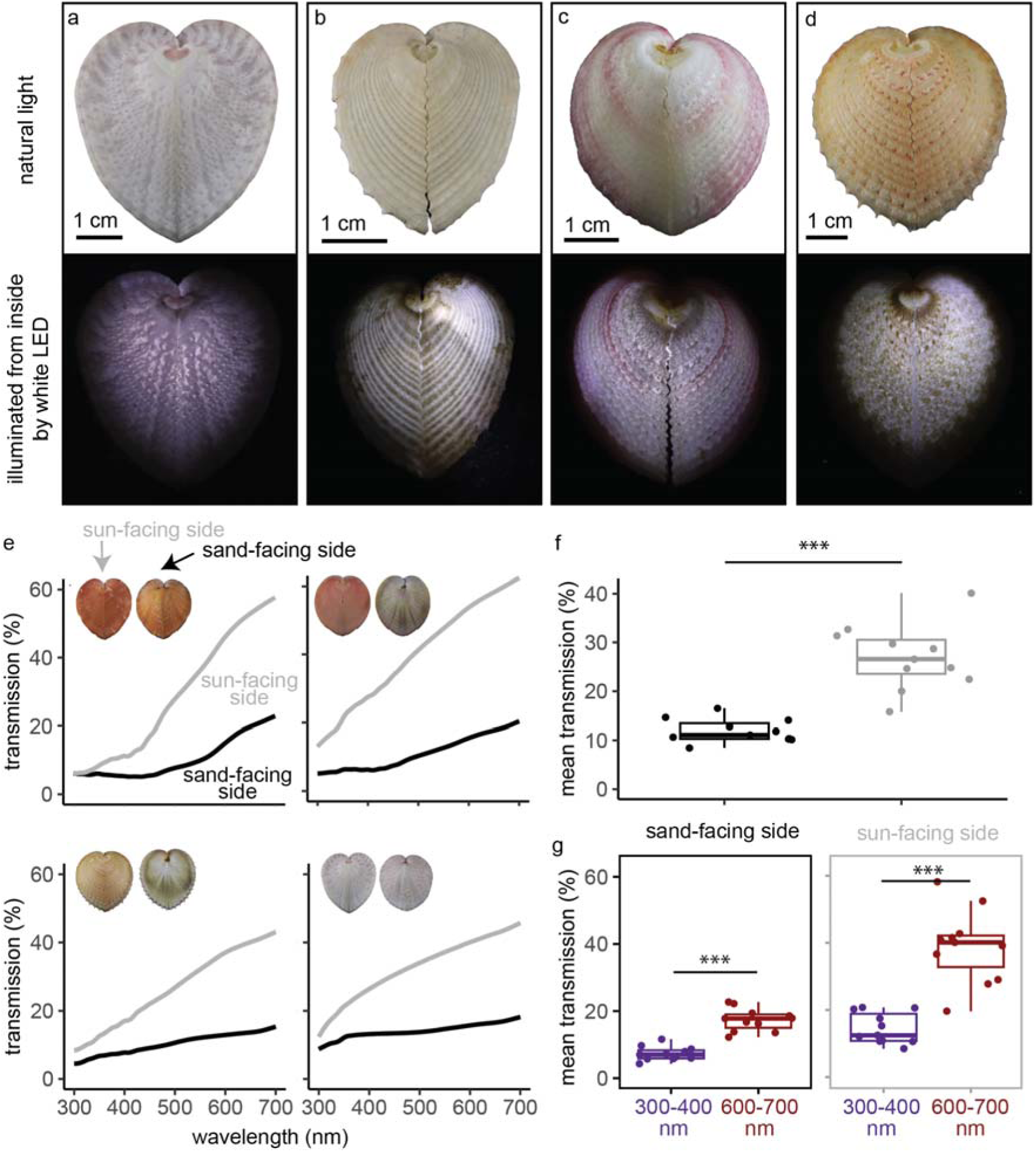
Transparent windows allow heart cockle shells to transmit 11-62% of photosynthetically active radiation (mean=31%) and significantly screen out UV radiation (mean=14%, range=5-28%). Here, we measure transmission through intact shells; in Supplementary Figure 1 we show that individual polished windows transmit 20-55% of light (mean=37%). **(a-d)** White LED lights inside each shell show transmission through the windows on the sun-facing side of the shell. Top row: natural light images; bottom row: lit up from inside. (a) *Corculum lorenzi*; (b-d) *Corculum cardissa.* (e) The shells’ transmission spectra rise at higher wavelengths for both the sun- and sand-facing sides (grey and black curves, respectively). Here we plot transmission for fourshells; see Supplementary Figure 2 for full results. **(f)** Averaging across 300-700nm, the sun-facing side of each shell has significantly higher mean transmission than the sand-facing side (two-sample two-sided paired t-test; p=<0.0005, 95% CI=[11, 19.2], mean diff.=15.1, df=10, t=8.22, Cohen’s *d=*3.03). **(g)** Shells transmit significantly more long-wavelength red light (600-700nm) than short-wavelength UV radiation (300-400nm), particularly on the sun-facing side (two-sample two-sided paired t-tests: sun-facing side; p=0.000012, 95% CI=[17.6, 31.3], mean diff.=24.4, df=10, t=7.98, Cohen’s *d*=2.93; sand-facing side: p=0.000019, 95% CI=[7.79, 12.4], mean diff.=10.1, df=10, t=9.8, Cohen’s *d*=3.60). The sun-facing side screens out UV radiation to a significantly greater extent than does the sand-facing side (two-sample two-sided paired t-test: p=0.00035, 95% CI=[8.31, 20.4], mean diff.=14.4, df=10, t=5.29, Cohen’s d=1.90). Boxplots show median, quartile 1, quartile 3, and whiskers that extend to the largest and smallest values ≤ |1.5 * interquartile range|. The sample size in **(f-g)** is n=11 distinct organisms. See Source Data for Figure 2 in UVVisTransmission_Corculumcardissa_16Mar2022.csv (Supplementary Data 1.zip)

The sun-facing side of the shell transmits substantial photosynthetically-active radiation to the symbionts within. We measured the spectral transmittance of the shell fragments in seawater using a UV-VIS spectrophotometer equipped with an integrating sphere to capture light transmitted at all angles. Across n=11 specimens, the sun-facing side transmits 11-62% of 400-700nm photosynthetically active radiation (mean=31%) but only transmits 5-28% of 300-400nm UV radiation (mean=14%; Figure 2e-f). In contrast, transmission through the sand-facing side is significantly lower. The sand-facing side transmits 4-25% of photosynthetically active radiation (mean=13%) and 2-13% of UV radiation (mean=7%; Figure 2e-f). Individual polished shell windows transmit 20%-55% of photosynthetically-active radiation (mean=37%, n=2; Supplemental Figure 1).

Further, the windows in the shell screen out UV radiation. Two to six times as much long-wavelength red light penetrates the sun-facing shell compared to UV radiation (comparing mean transmission from 600-700nm vs. 300-400nm; mean ratio=2.9, range of ratios=(1.8 - 6.2)), a significant difference (Figure 2g). While the sand-facing side of the shell also screens out UV radiation, the sun-facing side of the shell does so to a significantly greater extent (Figure 2g). The sun-facing side transmits 19-62% of 600-700nm red light (mean=38%). It is important to note that the wavelength range of oceanic illumination shifts toward blue-green colors with depth; transmission in the wild depends on this illumination as well as the optical properties of the shell.

Shells absorb more UV radiation than photosynthetically-active radiation and absorb almost no long-wavelength red light (Supplementary Figure 3). The sun-facing side absorbs, on average, 28% of UV radiation (range=5-68%) compared to only 6% of photosynthetically-active radiation (range=0-48%). The sand-facing side absorbs, on average, 40% of UV radiation (range=7-99%) compared to only 13% of photosynthetically-active radiation (range=0-77%). Shells absorb almost no 600-700nm red light, averaging 0% sunside and 2% sandside. Giant clams, members of the same family as the heart cockles and fellow photosymbionts, also screen or transform UV radiation through selective reflection as well as absorption and fluorescence ^8–10^.

We propose that the heart cockle’s ability to screen out UV radiation may be a protective adaptation to resist DNA damage and reduce bleaching risk from high-energy UV radiation. Light stress, and particularly UV radiation, can cause DNA damage, bleaching, and other problems for marine organisms ^25–29^. Other photosymbiotic animals, such as corals and giant clams, expose their soft tissues and symbionts directly to sunlight and UV radiation. Shell windows are an adaptation that allow the shallow-living heart cockles to screen solar radiation in a wavelength-dependent manner, due to a combination of at least two physical processes: CaCO_3_ in bivalve shells has strong absorption in the UV range, and scattering that tends to be inversely proportional to wavelength based on nano-scale inclusions in the shell and the intrinsic birefringence of CaCO_3_^30,31^.

### Condensing lenses beneath each window focus sunlight

We found that some *C. cardissa* individuals have small transparent truncated bumps on the interior of their shells, located exclusively beneath each window; our simulations show that these bumps act as simple condensing lenses (Figure 3). To obtain the precise 3D surface morphology of these bumps without damaging the specimens (i.e., with no contact), we used a laser scanning microscope which applies confocal scanning, focus variation, and white light interferometry to obtain their 3D geometry. We centered the microscope on individual triangular windows. The triangular perimeters of each bump consistently matched the roughly triangular shape of the windows (see three sample bumps in Figure 3c-e), but bumps varied in diameter and height (Figure 3c-h). We imported the 3D morphology of the interior bumps (Figure 3c-h) into optical simulation software to test how the bumps affect light penetration into the soft tissues of the heart cockle (schematic in Figure 3a-b).

**Figure 3:**
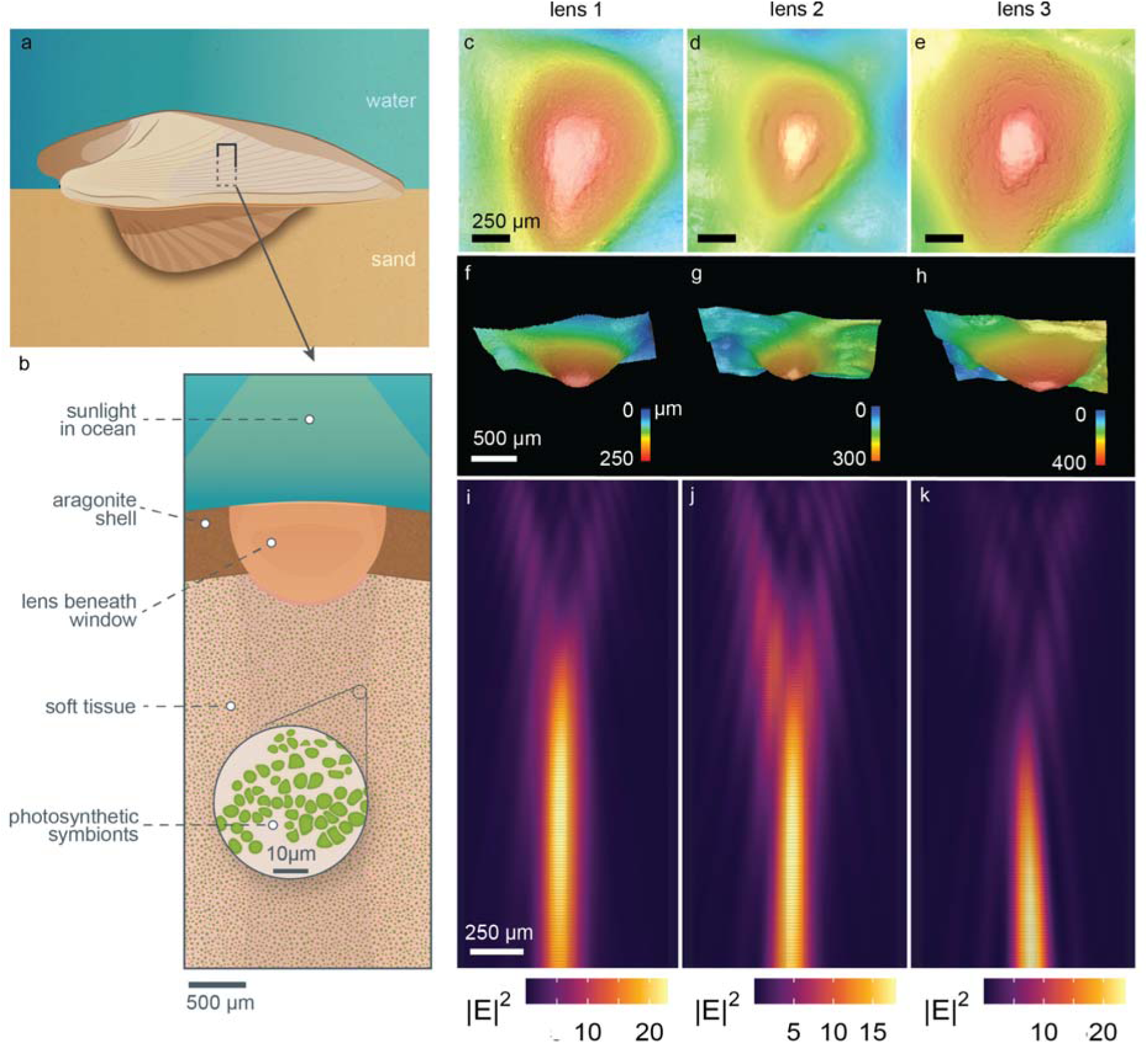
Condensing lenses with truncated caps under the windows focus sunlight. By “truncated”, we refer to the flattened tips of the lens. **(a-b)** A cross section of a heart cockle shell shows that beneath each shell window in some individuals, the aragonite forms a lens shape roughly 750μm in diameter. The exact size and shape of the lens varies from shell to shell. Beneath the shell is soft tissue with photosynthetic symbionts. **(c-e)** 3D images of three lenses viewed from below and **(f-h)** viewed from the side show truncated tops and variation in size and shape. **(i-k)** We imported the 3D scans in (c-h) into finite-difference time-domain (FDTD) optical simulation software to demonstrate that the lens focuses sunlight into a parallel beam. The input light geometry is a broadband (400-700nm) plane wave. The wavelength of light pictured in i-k is 625nm; lensing for other wavelengths of light, and with an extended vertical-axis, is shown in Supplementary Figure 5. We validated these simulations with lensing experiments using photosensitive paper, presented in Supplementary Figure 4. Illustrations in (a-b) are credited to Nuria Melisa Morales Garcia. Source Data for Figure 3 can be found in lensing_efield_results.zip (Supplementary Data 1.zip)

Our simulations demonstrate that the truncated lenses condense light with a ∼1mm depth of focus beginning around 500-750μm below the bump (Figure 3i-k). We propose that the bumps are an adaptation to help sunlight penetrate more deeply into symbiont-rich tissues ^12^. Many zooxanthellae are concentrated beneath the surface in the delicate, thin mantle tissue and gill filaments, suggesting that the lenses’ depth of focus may correspond to symbiont location^12,15^. To experimentally validate the lensing simulations, we recorded qualitative lensing behavior by placing shell fragments at different heights above photosensitive paper in natural sunlight (Supplementary Figure 4). Previous researchers proposed that these bumps either disperse light over interior tissues (e.g., ^23,24^) or condense light into a beam that can penetrate deeper into the zooxanthellae-rich tissue (e.g., ^15^). Our results support the hypothesis that the lenses condense light. Acetate peels from past work show that the aragonite microstructure within the bumps is a typical dissected crossed prismatic structure (rather than specialized fibrous prisms; see following section^15^).

Microlenses are widespread in nature as adaptations to manipulate light^32^. Here, microlenses may have evolved for photosynthesis by the symbionts in heart cockles; shade-dwelling angiosperm plants convergently evolved microlenses for photosynthesis through conical epidermal cells that concentrate sunlight ^33–35^ . Only some heart cockle individuals have lenses, suggesting that there is more to the evolutionary story than we currently know. Heart cockle microlenses are made of aragonite; similarly, chiton molluscs use aragonite lenses for vision ^36^ while certain other marine creatures see or sense light with calcite lenses, for example, in light-sensitive brittlestar arms ^37^ and the schizochroal eyes of certain extinct trilobites ^38^. Microlenses in nature can also produce richer colors by concentrating light onto pigments and reducing surface reflectance, as in the conical cells of flower petals ^39–41^ and super black color in peacock spiders (*Maratus*, Salticidae) ^42^.

### Fiber optic cable bundles, composed of aragonite, form the microstructure of windows

In the shell windows, the mineral microstructure forms bundles of parallel natural fiber optic cables to transmit light (Figure 4). These “fibrous prismatic crystals” are elongated spires oriented roughly perpendicular to the shell surface (as revealed by SEM, Figure 4a-b; see also microscopy in Supplementary Figure 6). Fiber optic cable bundles (also termed “mosaics” in their rigid form) are bound, co-aligned fibers that can – if the fibers are co-terminal– transmit images ^43^. Each shell window is itself a fiber optic cable bundle. The aragonite fiber optic cable bundles extend through an average of 83% of the shell thickness (range=70-99%; n=9 polished cross sections of windows; see Supplementary Figure 6). The fibers either terminate at a crossed lamellar portion of aragonite or the outer protective periostracum layer of the shell.

**Figure 4:**
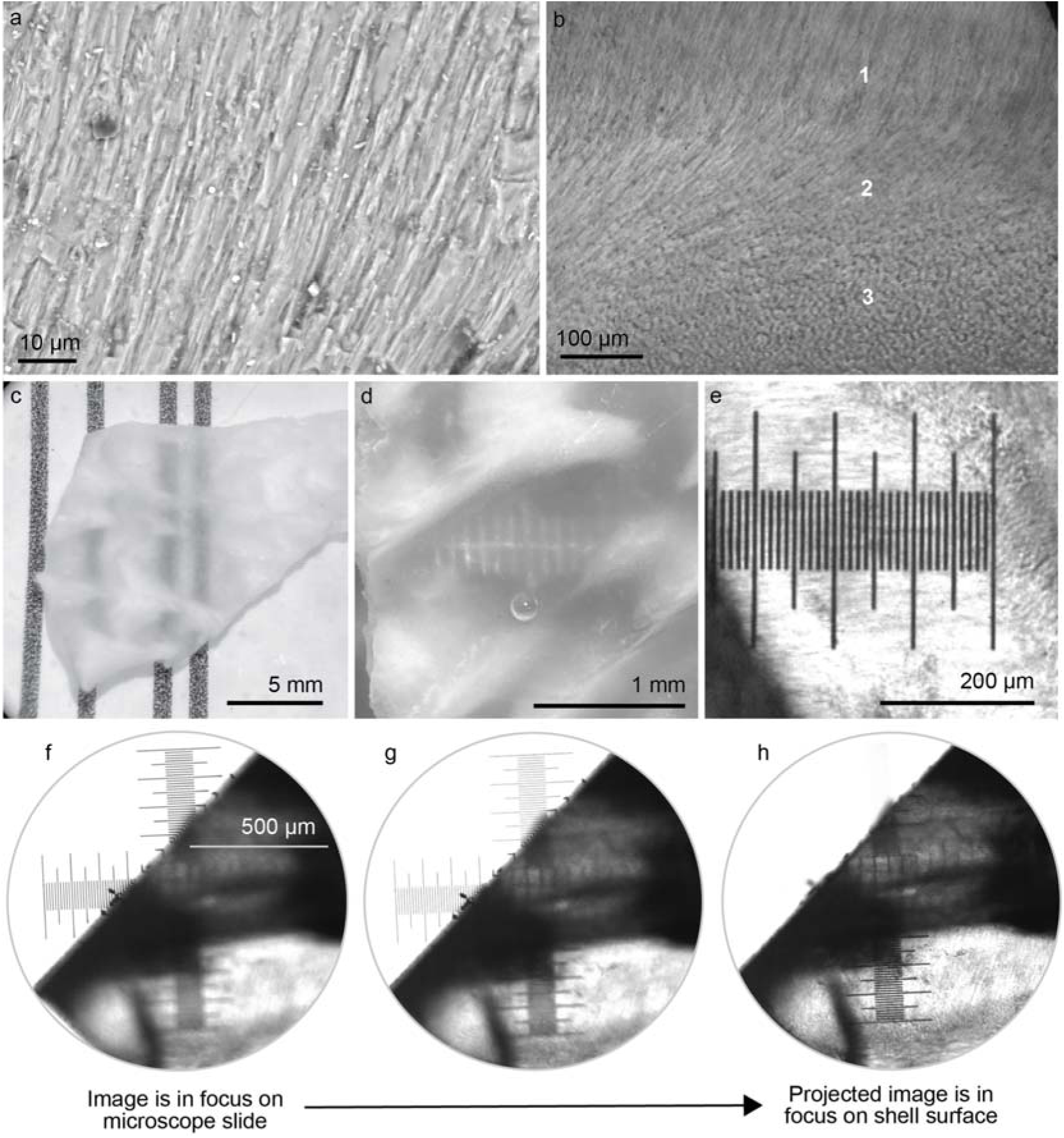
Shell windows use natural fiber optics to project images with very high resolution (100 lines/mm). **(a)** A cross-sectional SEM of a single shell window shows fibrous prisms of aragonite oriented roughly perpendicularly to the shell surface (the “fibrous prismatic layer”). Together, these prisms act like fiber optic cable bundles. **(b)** A light microscope image of a polished shell fragment shows the fibrous prisms rotating in orientation between the regions labeled 1-3. In region 1, we see the side view of fibrous prisms; in region 3 we see cross sections of aligned prisms pointing into the page. **(c-e)** Shell windows transmit high resolution images. **(c)** A fragment of unpolished shell shows the contrast between windows (transparent regions that transmit the vertical black lines) and the opaque surrounding shell (white regions that occlude the vertical black lines). **(d)** Shell windows from an unpolished fragment transmit 10 lines / mm. **(e)** Shell windows from a polished fragment transmit 100 lines / mm. **(f-h)** To experimentally test whether the windows act like fiber optic cable bundles (Supplementary Figure 7), we placed a small 0.3mm thick polished fragment of shell from a heart cockle on top of a glass calibration slide. We focused the microscope on the ruler on the glass calibration slide and then adjusted the focus to refocus on the surface of the shell (∼0.3mm higher than the background glass slide). By doing so, we could test whether images are *transmitted* through the windows or are actually *projected* onto the surface of the shell. Shell windows project images onto their surface (rather than merely transmitting images through). The image came into focus on the surface of the shell as we varied the plane of focus. Here we show images and micrographs from **(a,c,d)** two representative unpolished shell fragments and **(b,c,f,g,h)** one representative polished shell fragment. In **(c-d)**, shell fragments were immersed in seawater. Illustration in **(f)** is credited to Nuria Melisa Morales Garcia.

As stated above, the shells of heart cockles (and many other marine invertebrates) are made from aragonite ^19^, a crystal form of calcium carbonate CaCO_3_, inside an organic matrix of beta-chitin and other materials ^20,44,45^. In opaque regions of heart cockle shells, aragonite tends to be planar and crossed in orientation (termed “crossed lamellar”, either branching, complex, simple, or cone-complex ^15,16^)-- a common morphology that makes shells harder to break but also opaque.

Further, aragonite is orthorhombic, and the aragonite fibers in heart cockle shell windows are co-oriented along the mineral’s c-axis (as revealed by FTIR; Figure 5). The c-axis has the highest refractive index of aragonite’s three optical axes: 1.530 (a-axis), 1.681 (b-axis), and 1.686 (c-axis) ^46^.

**Figure 5:**
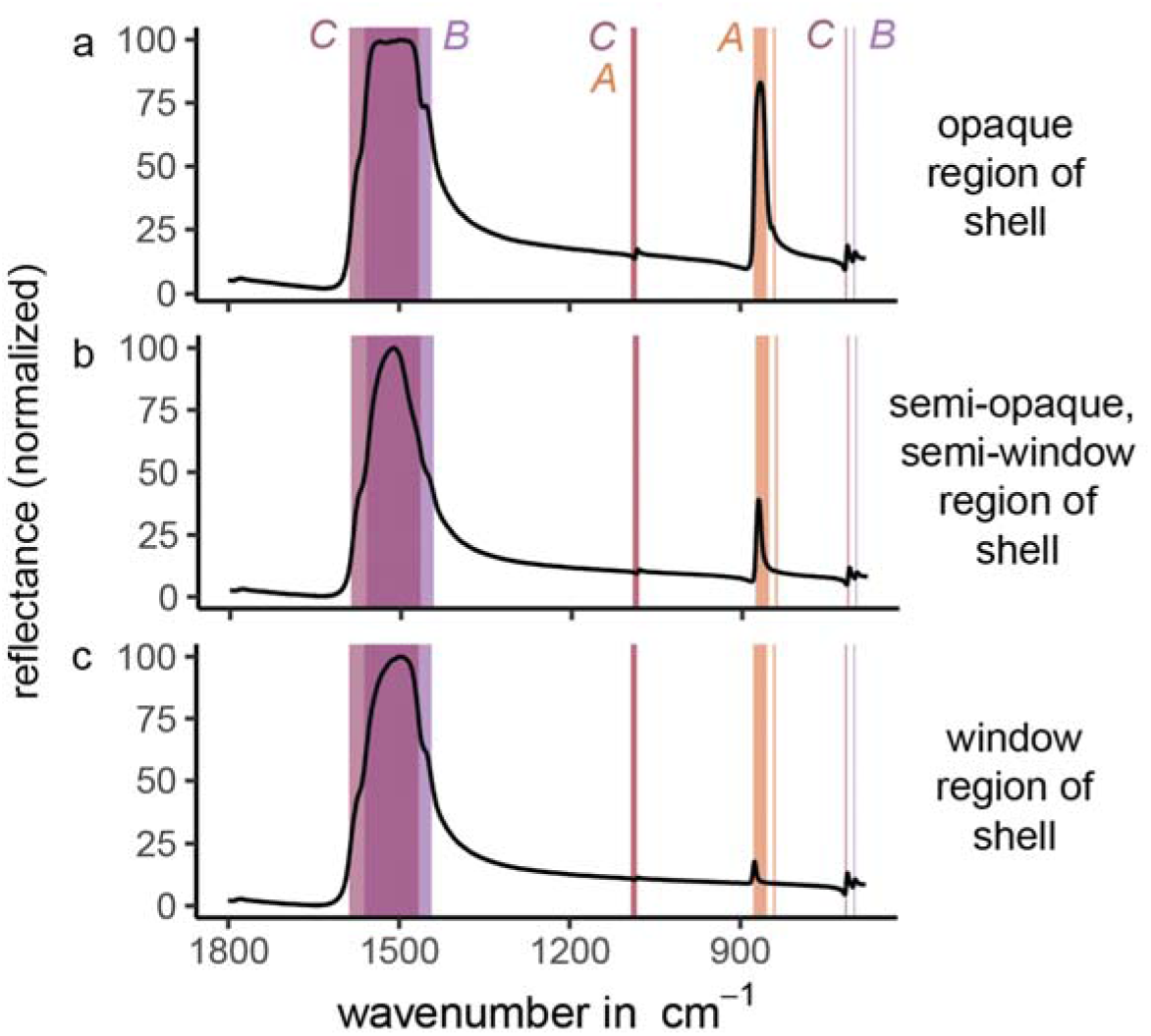
FTIR demonstrates that shell windows have a high proportion of c-axis aligned aragonite crystals compared to opaque regions. Here we plot three characteristic spectra taken from polished fragments of shell. Peak location corresponds to crystallographic orientation ^59,60^; a-axis orientation (“A” in figure) is highlighted in salmon, b-axis-orientation (“B”) in pale purple, and c-axis-orientation (“C”) in pink. B- and c-oriented crystals produce a very similar spectra, but the peak located between 853-877 cm^-1^ is unique to a-oriented crystals. **(a)** Reflectance from an opaque region of the shell shows a high proportion of a-oriented crystals. **(b)** Reflectance from a semi-opaque, semi-window region of the shell shows a medium proportion of a-oriented crystals. **(c)** Reflectance from a window region of the shell shows a low proportion of a-oriented crystals. Because the c- and b-axes of aragonite are so similar, it is difficult to distinguish which of these higher refractive indices is oriented in the Z direction. According to the literature, the c-axis is generally thought to be parallel to the long axis of aragonite prisms ^61^; therefore, we make the same conclusion. Whether it is the c-axis, b-axis, or both, our results do not materially change. Source Data for Figure 5 can be found in FTIRMeasurements_29April2022.zip (Supplementary Data 1.zip)

High-resolution images are visible through unpolished (Figure 4c-d) and polished (Figure 4e) fragments of shell windows. Unpolished windows transmit images at a resolution of 10 lines/mm (Figure 4c-d); polished windows transmit at 100 lines/mm through 300μm thickness (Figure 4e). Other researchers have observed these fibrous prismatic crystals of aragonite and debated whether or not they have a purpose, e.g., to act as fiber optic cables ^13–16,21,22^.

We show that the fibrous prismatic crystals act like parallel bundles of fiber optic cables in the shell windows– not just transmitting light but projecting high-resolution images through the window (Figure 4f-i, Supplementary Figure 7). An image in focus through a microscope is out of focus when a polished fragment of shell window is placed on top of it (Figure 4g); when the microscope is refocused (Figure 4h) onto the top plane of the polished shell window, the object returns into focus (Figure 4i). We observed the same phenomenon when marking the top of a shell window with a permanent marker dot, which then appeared to be projected onto the opposite surface of the shell. Heart cockle shell windows are therefore analogous to ulexite (the “TV stone”), which is composed of self-cladding fiber optic crystals that transmit images through fragments that are several cm thick ^47,48^. Heart cockle shell windows project images at a higher resolution (∼100 lines / mm; Figure 4e) than ulexite (∼10lines/mm ^47^), likely because the individual fibers are narrower and the shells are thinner. It is not clear that projecting images is of any use to the heart cockle; perhaps the lenses are associated with sensory perception in addition to photosynthesis. However, these experimental results show that the mineral is acting as a bundle of fiber optic cables.

To our knowledge, heart cockle shells are the first example of fiber optic cable bundles in a living creature. Indeed, fiber optic cables are themselves rare in nature. Certain deep-sea hyperiid amphipods have crystalline cones which act as light guides via fiber optic principles (e.g., ^49,50^). *Phronima sp.* looks upward at downwelling light to catch prey; they have tapered cone cells about 1 mm long and up to 185 micrometers in diameter that guide and focus light to enable accurate vision in low-light conditions ^49^. The amphipod *Hyperia galba* has tapered cone cells about 300-600μm long and 45-85μm wide, with graded refractive indices to focus light and screen off-axis light– an arrangement which allows the amphipod to see while remaining transparent ^50^.

Beyond amphipod vision, two species of deep-sea sponge have glass spicules with fiber optical properties, although it is not known whether the sponges’ optical properties are useful to the animal or are a side effect of selection for mechanical rigidity. The Venus Flower Basket sponge *Euplectella aspergillum* grows an intricate cage out of narrow spicules of amorphous, hydrated silica which can function as optical fibers and show remarkable mechanical strength compared to human-made glass ^51–53^. The large Antarctic sponge *Rossella racovitzae* has flexible spicules of amorphous hydrated silica that conduct light, even across a 90° bend, and– intriguingly– may support photosynthesis in shade-adapted diatoms that adhere to the spicules ^54^. These creatures tend to be found in the deep sea, although their ranges can extend into the photic zone; *Rossella racovitzae* lives at depths ranging from 18 to 2000m ^55^, while *Euplectella* lives at depths between 35 and 5000m ^56–58^.

Human-made fiber optic cables do not transmit all wavelengths of light with equal efficiency; low wavelength light scatters more due to imperfections and therefore is transmitted in lower proportions. The same phenomenon seems to occur in these natural fiber optic cable bundles, allowing heart cockles to protect their symbionts from UV radiation using the fiber optic structure of their shell windows. In heart cockles, the windows seem to screen UV radiation through two mechanisms: first, scattering due to imperfections in the fiber optic structures and CaCO_3_‘s birefringence, and second, wavelength-specific absorption from CaCO_3_ and yellow/orange shell pigments (Supplementary Figure 6) ^30,31^.

### Computational parameter sweeps show that the fiber optic aragonite morphology, size, and orientation transmit more light than many other possible arrangements

We conducted a series of computational parameter sweeps to demonstrate that the observed morphology of the aragonite fibers transmits more light than many other possible morphologies. Specifically, we performed finite-difference time-domain (FDTD) and finite element method (FEM) simulations using numerical optical solvers; we varied the morphology, size, and orientation of the fibers and extracted the transmittance from the simulations. The simulated fibers transmitted the most light when they matched our experimentally-observed morphology (consisting of fibrous prisms rather than lamellar planes; Figure 6a-c), size (width around 1μm; Figure 6d), and orientation (along the c-axis; Figure e-f). We validated these simulations by experimentally measuring transmission through polished shell windows (Supplementary Figure 1). Indeed, simulations at the observed parameter values matched well with experimental measurements showing that 300µm-thick windows transmit 32%-46% of 400-700nm light (gray rectangles in Figure 6A,D,E).

**Figure 6:**
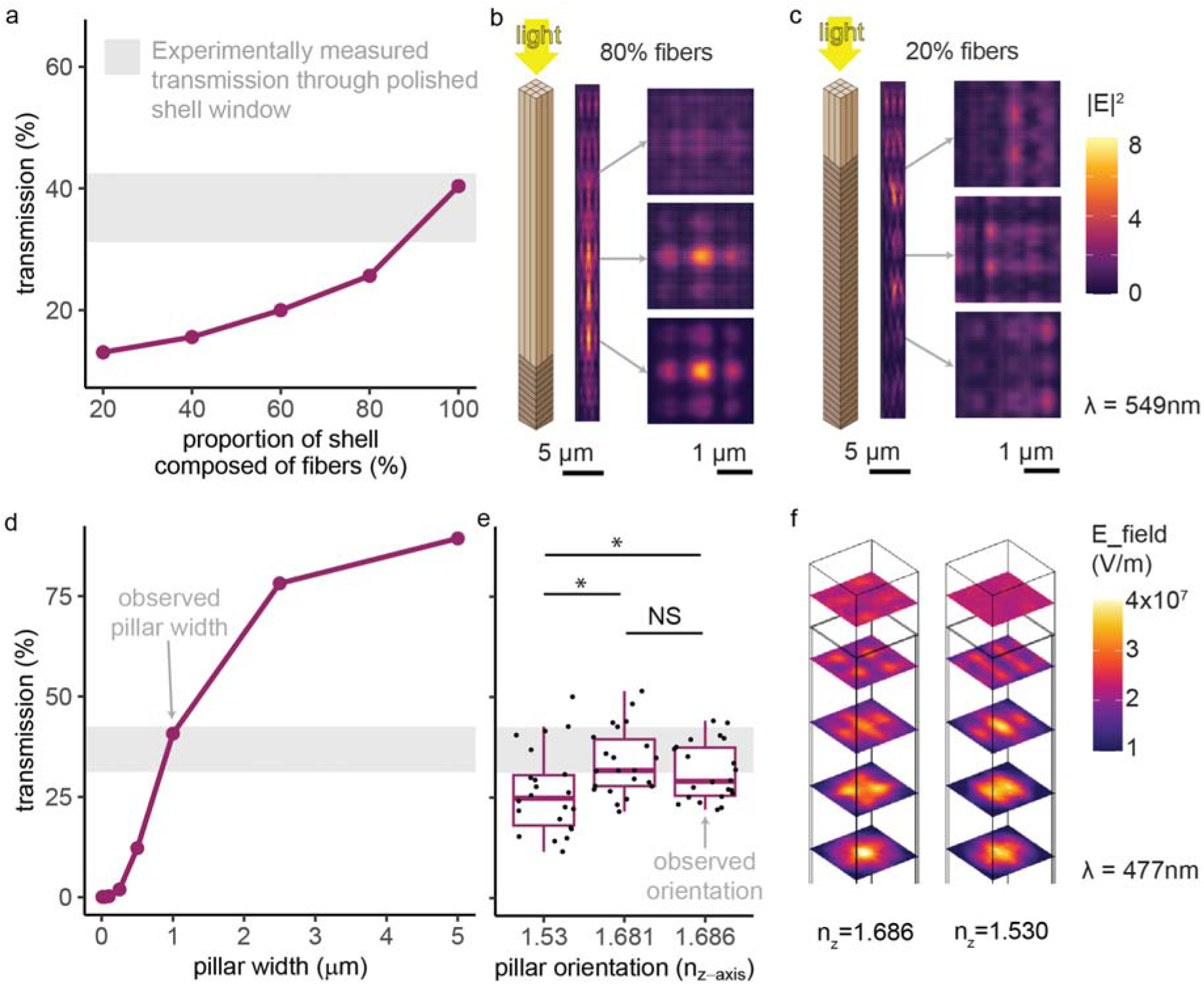
The observed orientation, size, and shape of the aragonite crystals allow them to transmit more photosynthetically-active radiation (400-700nm) than other morphologies. We conducted parameter sweeps in simulation and validated the simulations by measuring transmission of 400-700nm light through 300μm-thick polished shell windows (Supplementary Figure 1; gray rectangle in **(a)**, **(d)**, and **(e)** = mean experimental transmission +/-1 sd). **(a)** More light is transmitted through shell windows with a higher proportion of fibers rather than planes. Simulations of 100% fibers match experimental measurements (gray rectangle). **(b-c)** Light propagates differently through shells composed primarily of fiber-optic-cable shaped (left) compared to planar (right) aragonite. **(d)** Transmission increases with fiber width. Simulations of the observed 1μm width match the experimental measurements (gray rectangle). **(e)** Orientation along one of the two higher refractive indices (b or c), the observed arrangement (Figure 5), transmitted significantly more light than orientation along the a-axis (two-sample two-sided paired t-tests: a vs. c, p=0.00092, 95% CI=[2.26, 7.55], mean diff.=4.91, df=21, t=3.85, Cohen’s *d=*0.55; a vs b, p=0.00058, 95% CI=[3.62, 11.3], mean diff.=7.44, df=21, t=4.05, Cohen’s *d*=0.79; b vs c, no significant difference). Boxplots show median, quartile 1, quartile 3, and whiskers that extend to the largest and smallest values ≤ |1.5 * interquartile range|. N=22 wavelengths were simulated. **(f)** Light propagation depends on a fiber’s optical orientation (for both fibers, n_y_=1.681; left, n_x_=1.530; right, n_x_=1.686). We conducted FDTD simulations in Ansys Lumerical for **(a-d)** and FEM simulations in COMSOL for **(e-f)**. Results were consistent when we varied the width and complex refractive index of the organic matrix (see Supplementary Figure 8 and Methods). Illustrations in **(b-c)** by Nuria Melisa Morales Garcia. See Source Data for Figure 6 in Lumerical_planar_vs_columnar_simulations_V2.csv, COMSOL_VaryRefractiveIndex_Simulation_Results_V11.csv, Lumerical_vary_pillarwidth_simulations_V1.csv, planarvscolumnar.zip (Supplementary Data 1.zip)

Fibrous prisms of aragonite transmit more light than traditional planar, lamellar shapes (Figure 6a-c). Optical simulations demonstrate that the greater the proportion of fibrous prisms versus plates (Figure 6b-c), the greater the light transmission through the shell (Figure 6a). This mechanism of transparency through fiber-optics contrasts with another transparent bivalve, the window-pane oyster *Placuna placenta*. The window-pane oyster (used by some as a substitute for glass ^62^), achieves transparency alongside mechanical strength through crystallographically co-oriented calcium carbonate plates oriented in parallel to the surface of the shell ^31^.

The ∼1μm diameter of the observed fibers is superior to smaller fiber diameters for light transmission, but inferior to all larger fiber diameters (Figure 6d). We speculate that the aragonite fibers evolved to that width as a compromise between light transmission and mechanical toughness. In its fibrous prismatic form, aragonite varies among marine bivalves in width (roughly, 0.5-5μm) and cross-sectional shape (researchers have observed rectangular, lath-type, rod-shaped, anvil-shaped, and irregular cross sections ^63,64^).

In shell windows, the aragonite fibers were oriented along their c-axis, the axis with the highest refractive index. Fiber optic cables transmit light through total internal reflection due to cladding with a material of a lower refractive index ^43^. Through a series of optical simulations, we showed that the c-axis orientation achieves significantly greater total light transmission compared to orienting along the lowest refractive index (Figure 6e-f), likely due to the larger difference in refractive index between c-axis aragonite “core” (n_c_=1.686) and the “cladding” (a- and b-axes, n_a_=1.530, n_b_=1.681, and organic matrix,nm_atrix_=1.435). Indeed, the “TV stone” ulexite is self-cladded by its own anisotropy, with the center being oriented along the highest of its three refractive indices ^47^ – just like the aragonite reported here.

By orienting its fibers along the highest refractive index, the aragonite maximizes the “self-cladding” contribution from its own refractive indices ^47^ --because light traveling at an angle will encounter lower refractive indices--as well maximizing the traditional cladding from the organic matrix. Here, natural selection acted upon an intrinsic feature of aragonite, anisotropy, to produce a biologically useful outcome: more transmitted light for photosynthesis.

Another marine animal exploits the anisotropy of aragonite for vision. The chiton *Acanthopleura granulata* (Mollusca) lives on rocky substrates along seashores which are exposed to water during high tides and air during low tides; therefore, its eyes must be able to resolve images amidst background refractive indices of both air (n=1) and water (n=1.33). This chiton has aragonite lenses in its eyes which successfully focus light in both air and water thanks to the two different refractive indices of aragonite n_a_=1.53 and n_b_≈n_c_≈1.68 ^36^. Within the lenses, grains of aragonite are co-oriented along the c-axis ^65^.

Therefore, the observed morphology of shell windows seems to sit at a rough optimum for transmitting light. The shell is not entirely composed of windows, though, because transparency often compromises mechanical toughness in living creatures ^65^. Heart cockle shell windows are arranged in radial stripes, mosaics, or spots (Figure 2)– likely a balance between enhancing photosynthesis and avoiding shell-cracking predators. Chitons face the same tradeoff in their aragonite-lensed eyes; indeed, they have optimized for both mechanical toughness and optical function, in part by embedding the lenses– mechanically weak spots– within grooves to improve toughness ^65^.

### Determining biological function from optical properties

Just because a natural material has certain optical properties does not mean that those optical properties serve a biological purpose to the organism. Marvelous photonic properties may occur as a side effect of selection for a different purpose, such as toughness or smoothness. For example, blind golden moles (Chrysochloridae) have iridescent green, purple, and golden hairs arising from thin-film interference ^66^. But the moles are blind burrowers. Color seems irrelevant to their lives. For the moles, iridescence is likely a side effect of evolution for hairs that allow the moles to move smoothly through, and keep clean in, dirt ^66^. Photonic properties may also evolve due to randomness, genetic drift, physical constraint, or evolutionary history rather than to serve a specific biological purpose.

In contrast, we propose that the optical structures in heart cockles serve a biological purpose: transmitting necessary sunlight to photosynthetic symbionts. Windows and photosymbiosis are both rare among bivalves, and heart cockles must transmit light to their symbionts in order to survive. The windows transmit significantly more light than typical shells due to specialized optical properties, as our experiments show (Figure 2). The sun-facing side of shells is significantly thinner than the sand-facing side (Supplementary Figure 9), but increased transmission cannot simply be attributed to an overall thinner shell for two reasons. First, the windows are no thinner than those on the adjacent opaque shell and are, in some cases, thicker. Second, shell thickness does not correlate with light transmission across different individuals (Supplementary Figure 9). Thickness is perhaps the most important parameter in protecting shells against predation^67^, and heart cockles seem to have struck a compromise by making the shell a little thinner but getting most of their transmission gains from the optics inside the shell.

Further, simulations indicate that the size, shape, and orientation of aragonite fibers sit at a rough evolutionary optimum for transmitting light (Figure 6). Transmission is higher when the aragonite is formed into fibers, when the fibers are co-aligned, and when the fibers are specifically co-aligned along the c-axis or b-axis rather than a-axis (Figure 6). These parameter sweeps provide indirect evidence of biologically-selected function. Therefore, the specific optical properties of the windows seem essential for transmitting light.

Calculations of real illumination conditions in the tropical oceans also support the idea that shell windows evolved aragonite fibers for a specific purpose: to allow for efficient photosynthesis. Members of Symbiodiniaceae photosynthesize with the highest efficiency around 100 µmol quanta m^−2^s^−1^, and little or no further efficiency is gained from irradiance levels beyond 300 µmol quanta m^−2^s^−1 8,68,69^. Consider typical mean irradiance of 410 μmol photons m^−2^s^−1^ at 1 m depth in Singapore^70^, habitat for heart cockles. By adding windows to their shell, heart cockles more than doubled their internal irradiance of photosynthetically active 400-700nm wavelengths from 53 to 127 μmol photons m^−2^s^−1^ (i.e., from 13% transmission in windowless shell to 31% transmission in windowed shell– see Figure 2– of 410 μmol photons m^−2^s^−1^), a range of illumination where those gains significantly improve photosynthetic efficiency. The same calculations apply across heart cockle habitats: irradiance in tropical reefs at depths where heart cockles typically live is on the order of 5-500 μmol photons m^−2^s^−1^, depending on depth and shade (e.g., in Okinawa, mean daily maximum irradiance at 3-5 m depth is 401.8 μmol photons m^−2^s^−1^ in exposed reefs and 4.8 μmol photons m^−2^s^−1^ in shaded reefs ^71^; in Singaporean fringing reefs irradiance levels increased from 10m depth to 1m depth, ranging from 26.3-451.9 μmol photons m^−2^s^−170^; in the North Atlantic Bermuda platform at 8−10 m depth, maximum daily irradiance peaked at 283.90 μmol photons ^72^). It is important to note that light levels at depth change with the weather, time of day, water clarity, and other parameters.

### Conclusion and future directions

In summary, heart cockles have evolved transparent windows in their shells with what is, to our knowledge, the first example of bundled fiber optic cables in a living creature. The aragonite fiber optics transmit light to the cockle’s photosynthetic symbionts and, apparently as a side effect, project high-resolution images. Beneath each shell, condensing lenses focus sunlight to penetrate the symbiont-rich tissues. Together, the biophotonic arrangement screens out UV radiation, potentially protecting against the risk of bleaching or UV damage to DNA and other biomolecules. It would be worthwhile to compare the radiation spectrum experienced by symbionts in heart cockles to that of reef-building corals and to examine the biological role of fluorescence across reef-dwelling creatures.

Beyond heart cockles and giant clams, bivalves may contain other potential models for understanding the evolution of photosymbiosis. Researchers have observed opportunistic symbioses between photosynthetic algae and many other lineages of bivalves ^7^, including freshwater mussels *Anodonta cygnaea* and *Unio pictorum* ^73^, the clam *Fluviolanatus subtortus* ^74^, the scallop *Placopecten magellanicus* ^75^, the cockle *Clinocardium nuttallii* ^76,77^, and likely others (see ^7^ for comprehensive review). These “symbioses” (excluding the obligate symbioses in heart cockles and giant clams) are not known to be mutually beneficial and, indeed, may be parasitic ^22^. One indirect measure of mutualism, rather than parasitism, may be biophotonic adaptations in the host to maximize light transfer and screen out UV radiation (as we see in the heart cockles).

The heart cockles’ fiber optic cables and microlenses may inspire optical technologies. Previously, the glass spicules of sponges inspired lightweight mechanical architectures ^78^, while microlenses in peacock spiders inspired antireflective polymer microarrays ^79^.

## Methods

### Ethics and Inclusion Statement

All people who contributed substantially to this manuscript were invited to join as co-authors. No ethics approval was required for this research. We obtained permission for destructive sampling of museum specimens.

### Specimens

We obtained shells identified as members of the genus *Corculum* from the Yale Peabody Museum of Natural History (catalog numbers YPM108178, YPM108179, YPM108180, YPM108181) and from Jean Pierre Barbier, topseashells.com (catalog numbers, TS185203, TS163469, TS179955, TS177080, TS196641, TS179954, TS181027, TS164335, TS163836, TS196638, TS164403, TS196643). Because the shells are dry, the organic matrix within may not perfectly represent the organic matrix in living or recently collected shells. To better assess shell morphology, we sent seven shells to be prepared as polished thin sections by Burnham Petrographics, https://www.burnhampetrographics.com/.

### Transmission, Absorption, and Microscopy

To measure light transmission through and absorption by the intact unpolished shells, we first cut 1 cm^2^ square fragments of each shell (one from the sun-facing side and one from the sand-facing side) using a Raytech Blazer four-inch lapidary saw. We suspended the shell fragments in seawater in cuvettes and used a Universal Measurement Spectrophotometer (UMS), Agilent Technologies Cary 7000 UV-VIS-NIR equipped with an integrating sphere. To measure percent light transmission, the cuvette was placed just outside the integrating sphere; to measure percent absorption, the cuvette was placed in the center of the integrating sphere. This method captures the most biologically relevant and realistic metrics, because it includes all angles of transmitted light, uses a fragment of actual shell including both windows and opaque regions, and measures light transmitted through the shell fragment suspended in seawater (therefore capturing the natural difference in refractive index between shell and medium). All measurements were normalized to a cuvette filled with seawater with no shell fragment. All measurements ranged from 300 to 700nm, which includes ultraviolet radiation. We also wanted to measure light transmission through polished fragments of isolated shell windows in order to ground-truth our simulations. Therefore, we obtained 300μm-thick polished shell windows by sending shell fragments to Brand Laser Optics (Newark, California, USA). We illuminated the windows with a diffuse light source, the Ocean Optics ISP-Ref integrating sphere equipped with a tungsten-halogen lamp, and collected light transmitted through the shell window with a fiber optic cable normally oriented to the shell and attached to an Ocean Optics Flame Spectrometer.

To visualize light transmission through windows in intact shells, we placed single LEDs on a wire inside closed shells.

To visualize image transmission through windows in an unpolished and polished shell fragment, we used a Leica DM4000 M light microscope. We placed the fragment in a dish atop a microscope calibration slide (which allowed us to identify the resolution of transmitted images).

To gather surface microstructure information, we used a Keyence VK-X250/260K 3D Laser Scanning Confocal Microscope.

To obtain aragonite fiber structural information with minimal damage to the specimens, shell fragments were imaged using a JEOL JSM-IT500HR environmental scanning electron microscope. Prior to imaging, the fragments were coated with ∼15nm of amorphous carbon to ensure electrical conductivity on the specimen surface.

### Raman and Fourier-Transform Infrared Spectroscopy

For Raman and infrared spectroscopy experiments, light dimension is reported in wavenumber (cm^-1^), which is conventional for vibrational spectroscopy. This value is defined as the number of lightwave oscillations per distance, and is the inverse of wavelength.

We confirmed that the shells, both windows and opaque regions, were composed of aragonite using Raman spectroscopy on the Horiba LabRAM HR Evolution Raman microscope and comparing the observed spectra to published literature ^19,80,81^. The shells were illuminated with a 532nm laser powered at ∼11 mW with a 600 l/mm grating with ∼1 cm^-1^ spectral resolution. Raman spectra were collected from 100 cm^-1^ to 1650 cm^-1^ using a 100x, 0.9 NA objective with a diffraction-limited spot size of ∼1.5μm and 15 second acquisition time.

To identify the crystallographic orientation of the aragonite in shell windows compared to opaque regions of the shell, we measured reflectivity using a Fourier transform infrared radiation spectroscopy (FTIR) setup. For these measurements, we collected the spectrum of infrared light (400 cm^-1^-1800 cm^-1^) reflected from shell windows using a Nicolet Continuum infrared microscope and a Nicolet iS50 FTIR spectrometer. These measurements were collected with 4 cm^-1^ resolution. The mirror in the interferometer moved at a velocity of 1.8988 cm/s when it scanned through the Fourier-transformed spectrum (slower optical velocities give more accurate measurements). All results were averaged over 100 spectra. We used polished samples of shell that included both opaque and transparent regions, and we measured at three points: (i) transparent window, (ii) at the border between window and opaque, and (iii) opaque. By comparing the resulting spectra to published literature ^59,60^, we were able to identify differences in crystallographic orientation between windows and opaque regions.

### Optical Simulations

We performed optical simulations using both idealized and actual, imported microstructures. We performed finite-difference time-domain (FDTD) simulations in the software program Ansys Lumerical and finite element method (FEM) simulations in the software COMSOL.

To test whether the truncated lenses focused or dispersed light, we used FDTD simulations based on measured 3D morphological scans. To save computer memory, we imported 3D surface files (STL triangular surface files) with a scaling factor of 0.0001 (one-tenth the actual size of the lenses). We launched a plane wave normally incident (z-direction) on the lens, ranging in wavelength from 400 to 700nm, and bounded the simulation on all sides by perfectly matched layers (PMLs). We placed frequency domain field monitors at regular intervals to identify the focal region of greatest intensity, then we centered XZ and YZ monitors on that focal spot.

To test the transmittance of aragonite in a fibrous versus a lamellar/planar shape, we performed FDTD simulations of a repairing element in the XY plane consisting of a 3 x 3 grid of fibers, each measuring 1 x 1 x 50μm. We varied the proportion of the 50µm stack that was composed of planar (rather than fibrous) aragonite. For the aragonite fibrous prisms, we used the refractive index values of 1.530 (a-axis), 1.681 (b-axis), and 1.69 (c-axis). For the organic matrix, we chose a real refractive index of 1.43 following past work and varied the imaginary component over two values: 0.0001 and 0.001 ^44,45^. We assume the organic matrix is not anisotropic. We report the results for imaginary index 0.001 in the main text and for 0.0001 in Supplementary Figure 8. Generally, molluscan shells are about 0.1-5% organic matrix by weight with the rest being calcium carbonate ^82^; we varied the width of the organic matrix between carbonate fibers over three values: 100nm, 50nm, 25nm ^83^. We report the results for organic matrix width 100nm in the main text and the others in Supplementary Figure 8. For simulations incorporating layers of planar aragonite, we incorporated planes with thickness 1μm in the Z direction and XY area spanning the entire XY plane of the simulation. The angle of the planes was randomly varied between 5 and 15 degrees (using the random() function). For these simulations, the simulation domain was bounded in the Z plane by perfectly matched layers and in the X and Y planes by periodic boundary conditions (to simulate an infinite array of fibers). We used a plane wave normally incident (z-direction), ranging in wavelength from 400 - 700nm.

To test the impact of varying the size of the optical fibers, we repeated the above procedure but varied the fiber width over the following values: 10nm, 25nm, 50nm, 100nm, 500nm, 1μm, 2.5μm, and 5μm.

To test the impact of varying the optical axis of the fibers, we performed finite element method (FEM) simulations in COMSOL on an idealized single fiber measuring 1 x 1 x 10μm. We enabled Floquet boundary conditions in the X and Y planes to simulate the periodic nature of the aragonite fibers with ports above and below in the Z plane. We performed parameters sweeps to vary the X-axis, Y-axis, and Z-axis refractive indices over all six possible configurations (that is, we set n_x_, n_y_, and n_z_ equal to 1.686, 1.681, and 1.530 but systematically varied which axis had which index). In the results, we condense these six possibilities into the three different Z-axis possibilities. We swept over wavelengths from 485nm to 725nm by 20nm, as well as 477 to 720nm by 27nm; wavelength values were chosen to avoid round numbers which may have created spurious resonances. Here we report the total transmission through the bottom of the fiber.

For all simulations on idealized structures (aragonite fiber optic cables), we eliminated spurious resonances by changing the size of the modeled features over three values – 0.975, 1, and 1.025 times the actual size– and averaging the results ^84^. To make all simulations comparable to experimental results from the 300μm-thick polished shell fragments, we adjusted the simulation results to represent transmission values for 300μm-high fibers. That is, when the simulation modeled 50μm-high fibers and gave the result T_50μm=_transmission(50μm-high fibers), we report T_50_ ^6^=T =transmission for 300μm-high fibers. For simulations modeling 10μm-high fibers which gave the result T_10μm_=transmission(10μm-high fibers), we report T_10_ ^30^=T =transmission for 300μm-high fibers

### Experimental Test of Light Focusing

To qualitatively determine whether the microlenses beneath each window focus light, we placed shell fragments atop light-sensitive cyanotype paper at varying heights ^37^: 0.5mm , 1.25mm, 2mm, and 4mm. We used spacers, cardstock, and construction tape to elevate the shell fragments to known heights. We placed the cyanotype papers in direct sun at noon on a cloud-free day in Sunnyvale, CA for 3 minutes and 30 seconds. We performed these exposures in air, and therefore, repeated our optical simulations of lens focusing in a background medium of air rather than seawater (see Methods: Optical Simulations).

## Data Availability

All data are available as supplemental information in **Supplementary Data 1.zip.** Within that zipped folder, Source Data for each figure can be found as follows:

● Source Data for Figure 2 can be found in UVVisTransmission_Corculumcardissa_16Mar2022.csv (Supplementary Data 1.zip)
● Source Data for Figure 3 can be found in lensing_efield_results.zip (Supplementary Data 1.zip)
● Source Data for Figure 5 can be found in FTIRMeasurements_29April2022.zip (Supplementary Data 1.zip)
● Source Data for Figure 6 can be found in Lumerical_planar_vs_columnar_simulations_V2.csv, COMSOL_VaryRefractiveIndex_Simulation_Results_V11.csv, Lumerical_vary_pillarwidth_simulations_V1.csv, planarvscolumnar.zip (Supplementary Data 1.zip)
● Source Data for Supplementary Figure 1 can be found in, Transmission_Polishedwindows_Corculum-cardissa_18Aug2023.csv (Supplementary Data 1.zip)
● Source Data for Supplementary Figure 2 can be found in UVVisTransmission_Corculumcardissa_16Mar2022.csv (Supplementary Data 1.zip)
● Source Data for Supplementary Figure 3 can be found in Corculum-cardissa_absorbance_7May2024.csv (Supplementary Data 1.zip)
● Source Data for Supplementary Figure 4 and Supplementary Figure 5 can be found in lensing_efield_results.zip (Supplementary Data 1.zip)
● Source Data for Supplementary Figure 8 can be found in COMSOL_VaryRefractiveIndex_Simulation_Results_V11.csv (Supplementary Data 1.zip)
● Source Data for Supplementary Figure 9 can be found in Corculum_specimens.csv (Supplementary Data 1.zip)

Original samples used for this work are available upon request to the lead author and specified by the catalog numbers listed in Corculum_specimens.csv (Supplementary Data 1.zip).

## Code Availability

All code is available as supplemental information in **Supplementary Data 1.zip.** Specifically, within that zipped file the code file **HeartCockles_AllAnalysis.R** was used to perform all analyses and plot all figures.

## Acknowledgements

D.E.M. is supported by the Stanford Science Fellowship and the NSF Postdoctoral Research Fellowships in Biology PRFB Program, grant 2109465. We also acknowledge support from the NSF Waterman Award, Grant Number 1933624, which covered the salaries of E.K., L.H., and J.D. B.O is supported by the Bio-X Bowes Graduate Student Fellowship. Part of this work was performed in the Stanford Microchemical Analysis Facility (MAF). The MAF is part of the Stanford Nano Shared Facility, which is supported by the National Science Foundation under award ECCS-2026822. We are grateful to members of the Johnsen Lab at Duke University and the Dionne Lab at Stanford University for help and feedback; in particular, we thank Jefferson Dixon, Briley Bourgeois, Alexander Al-Zubeidi, Halleh Balch, Harsha Eragamreddy, Kai Chang, Parivash Moradifar, and Ariel Stiber. We are grateful to Nuria Melisa Morales Garcia for artwork. We appreciate the help and advice from experts at the Yale Peabody Museum and the Harvard Museum of Comparative Zoology, including Jennifer Trimble, Gonzalo Giribet, Eric Lazo-Wasem, and Lourdes Rojas.

## Author Contributions Statement

DEM, JAD, and SJ conceived the project. DHB prepared samples. EK and DEM conducted simulations. LKH, BO, and DEM performed Raman Spectroscopy and FTIR. DEM performed spectrophotometry. DEM wrote the first draft of the paper and all authors reviewed, edited, and commented on the draft. JAD and SJ jointly supervised the project.

