## Supplementary Information for "Heart cockle shells transmit sunlight to photosymbiotic algae using bundled fiber optic cables and condensing lenses"

### Title

### Supplementary Figures

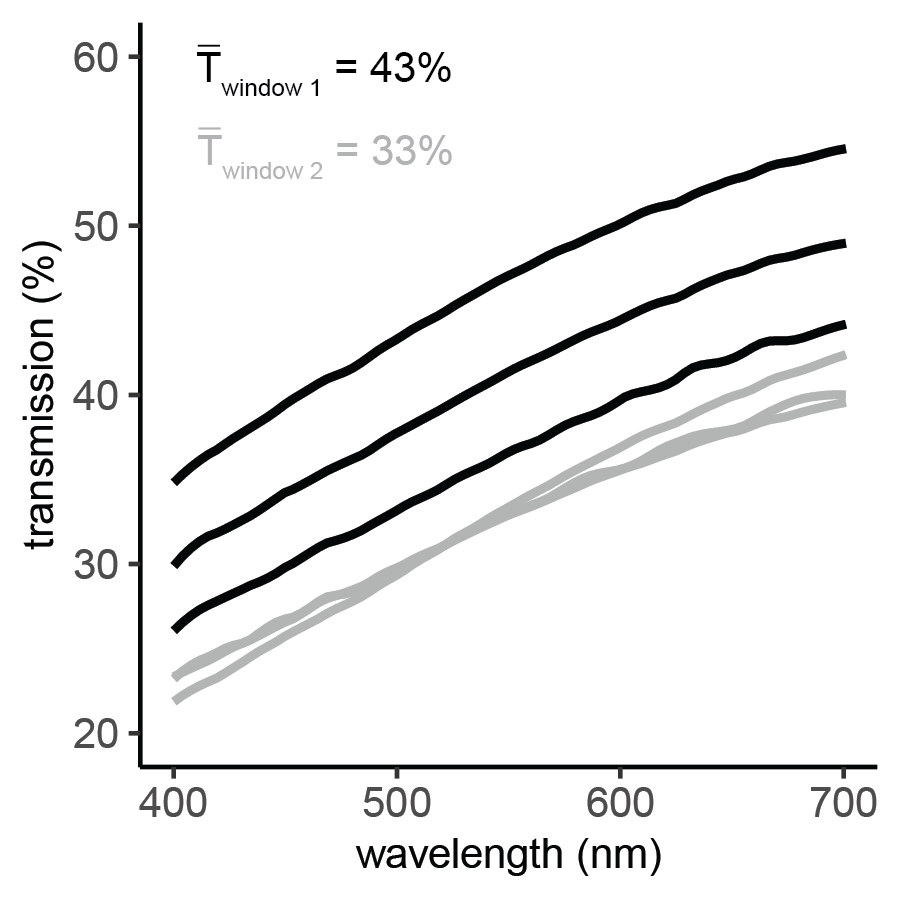

**Supplementary Figure 1. Individual polished shell windows transmit 20%-55% of photosynthetically-active radiation (mean = 37%).** We polished two individual shell windows to a width of 300 μm, so that we could ground-truth our simulations of light transmission (Main text, Figure 6). We measured transmission through the shell windows at three locations within each of two windows. Across 400-700nm, window 1 transmitted 24-55% (mean=41%) of light, while window 2 transmitted 20-43% (mean=33%). Source Data for Supplementary Figure 1 can be found in, Transmission_Polishedwindows_Corculum-cardissa_18Aug2023.csv (Supplementary Data 1.zip)

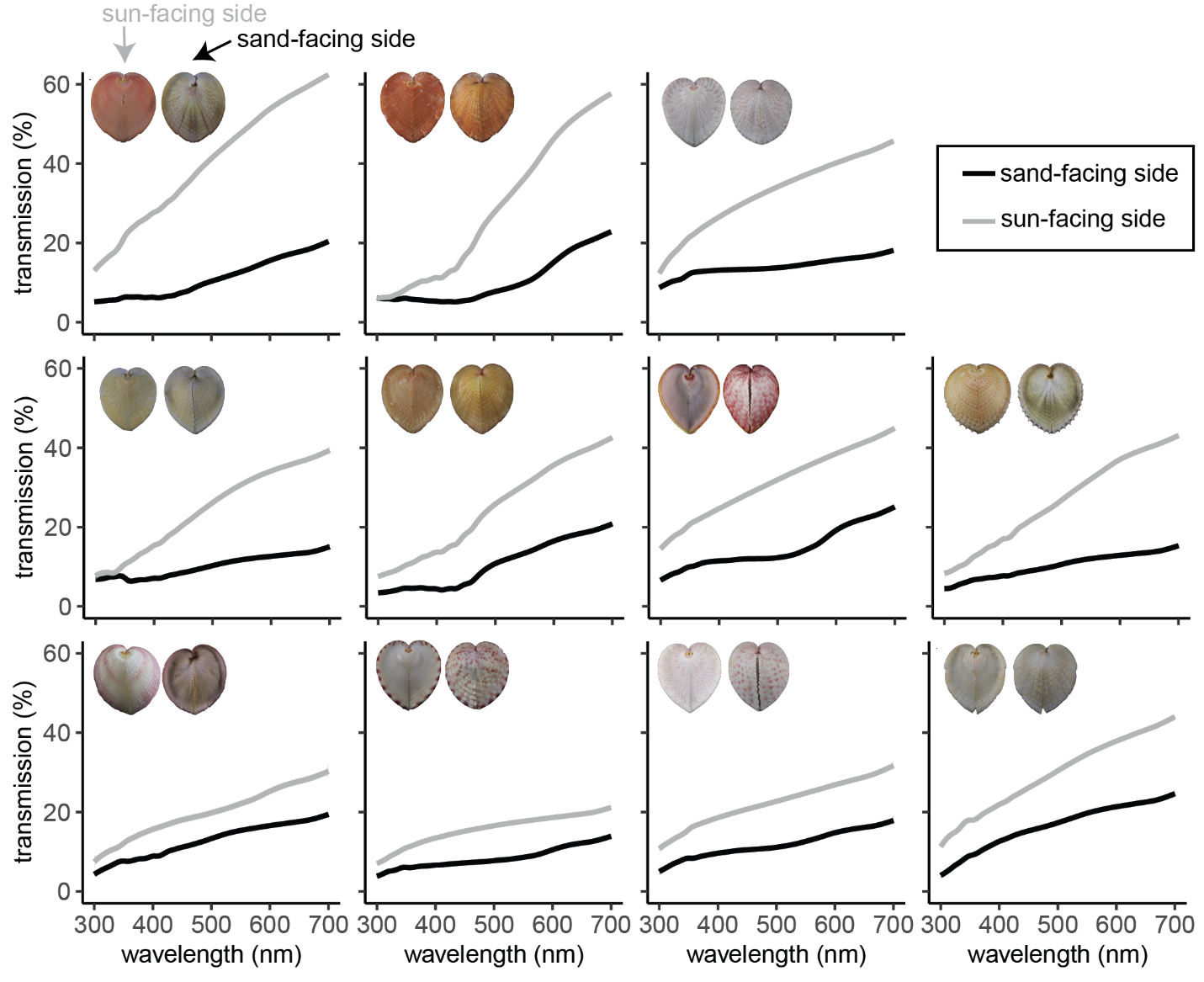

**Supplementary Figure 2: Transparent windows transmit 11-62% (mean = 31%) of photosynthetically active radiation, and 5-28% (mean = 14%) of UV radiation, into the soft tissues.** Gray lines indicate % transmission through the sun-facing (upper) half of the shell, and black lines indicate % transmission through the sand-facing (lower) half of the shell. The sun-facing side transmits 11-62% of 400-700nm photosynthetically active radiation (mean = 31%) but only transmits 5-28% of 300-400nm UV radiation (mean = 14%). The sand-facing side transmits 4-25% of photosynthetically active radiation (mean = 13%) and 2-13% of UV radiation (mean = 7%). Individual polished shell windows transmit 20%-55% of photosynthetically-active radiation (mean=37%, n=2; Supplementary Figure 1). For these measurements, a flat rectangular piece of shell approximately 1.5 cm by 1 cm was immersed in seawater. Each line represents three measurements, but error bars are small and concealed within the line width. Source Data for Supplementary Figure 2 can be found in UVVisTransmission_Corculumcardissa_16Mar2022.csv (Supplementary Data 1.zip)

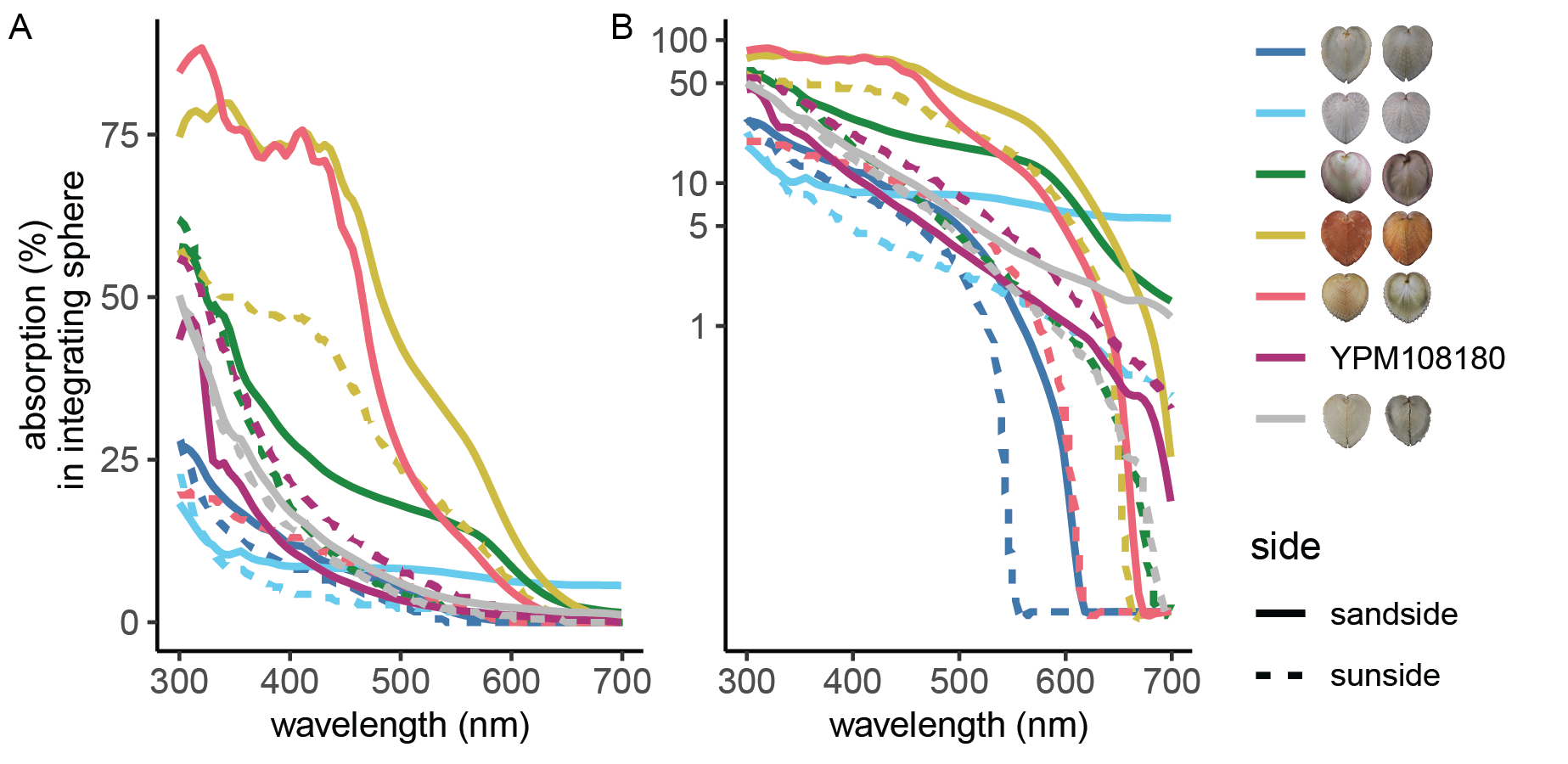

**Supplementary Figure 3: Heart cockle shells absorb light in a wavelength-specific manner.** We measured the absorption of n=7 shells. The same data are plotted here on **(A)** a normal scale and **(B)** a log scale. Shells with more orange/yellow pigment seemed to have greater wavelength-specific absorption (red-orange and yellow lines). For these measurements, a flat rectangular piece of shell approximately 1.5 cm by 1 cm was immersed in seawater in a cuvette. The cuvette was placed inside an integrating sphere and illuminated with normally-incident light. For specimen YPM 108180, no photograph was available for legend inset. Source Data for Supplementary Figure 3 can be found in Corculum-cardissa_absorbance_7May2024.csv (Supplementary Data 1.zip)

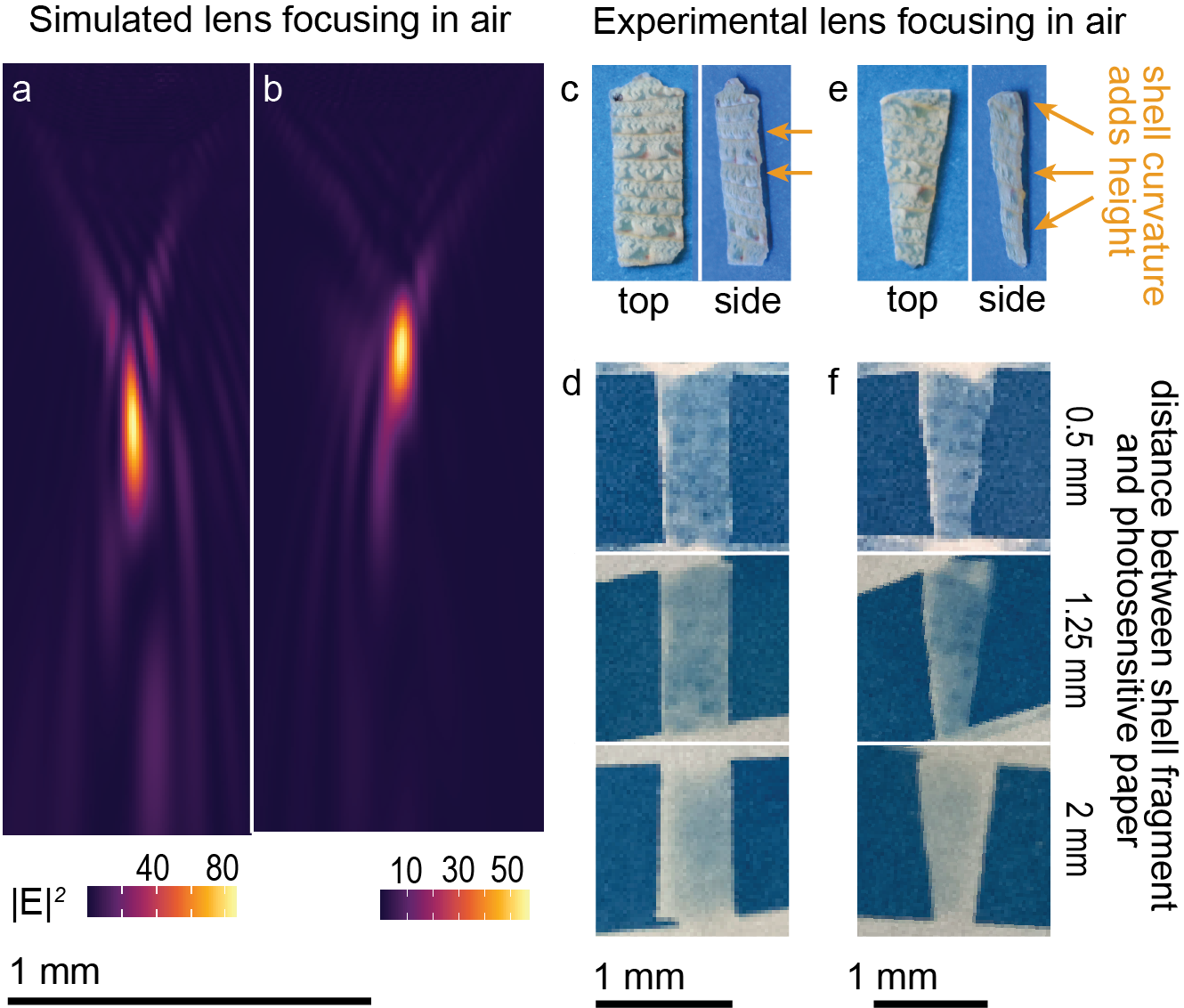

**Supplementary Figure 4. Microlenses focus light in qualitative, experimental validation of simulations.** We used photosensitive paper to create imprints of solar irradiation at different depths beneath shell fragments; these experiments were done in air using natural sunlight. **(a-b)** We repeated our Lumerical simulations of the lens focusing with air as the background medium. The most intense region of focus typically ranged from 0.5-1 mm below the shell surface. **(c-f)** We placed a shell fragment atop photosensitive cyanotype paper to visualize the lensing effect of light focusing and dispersing (**d** and **f**; top to bottom) at three depths: 0.5mm, 1.25mm, and 2 mm. However, these depths are not absolute, because the shell fragments had some natural curvature (**c**, **e**: orange arrows) that added roughly 0.25-0.75 mm of height. The depths reported in **(d-f)** were measured from the lowest points on the shell fragments. Source Data for Supplementary Figure 4 can be found in lensing_efield_results.zip (Supplementary Data 1.zip)

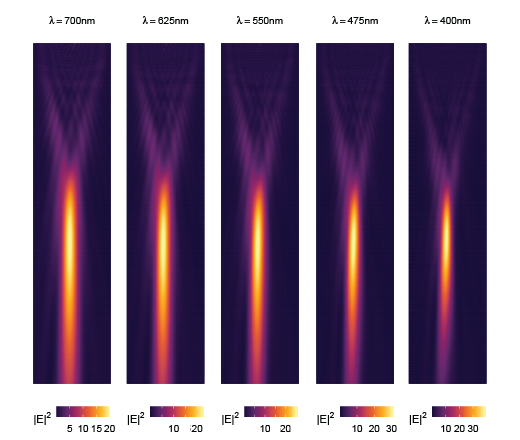

**Supplementary Figure 5: Chromatic aberration in condensing lenses beneath shell windows.** The lenses beneath shell windows (found in some individuals; see Figure 3) focus light differently depending on the wavelength. Source Data for Supplementary Figure 5 can be found in lensing_efield_results.zip (Supplementary Data 1.zip)

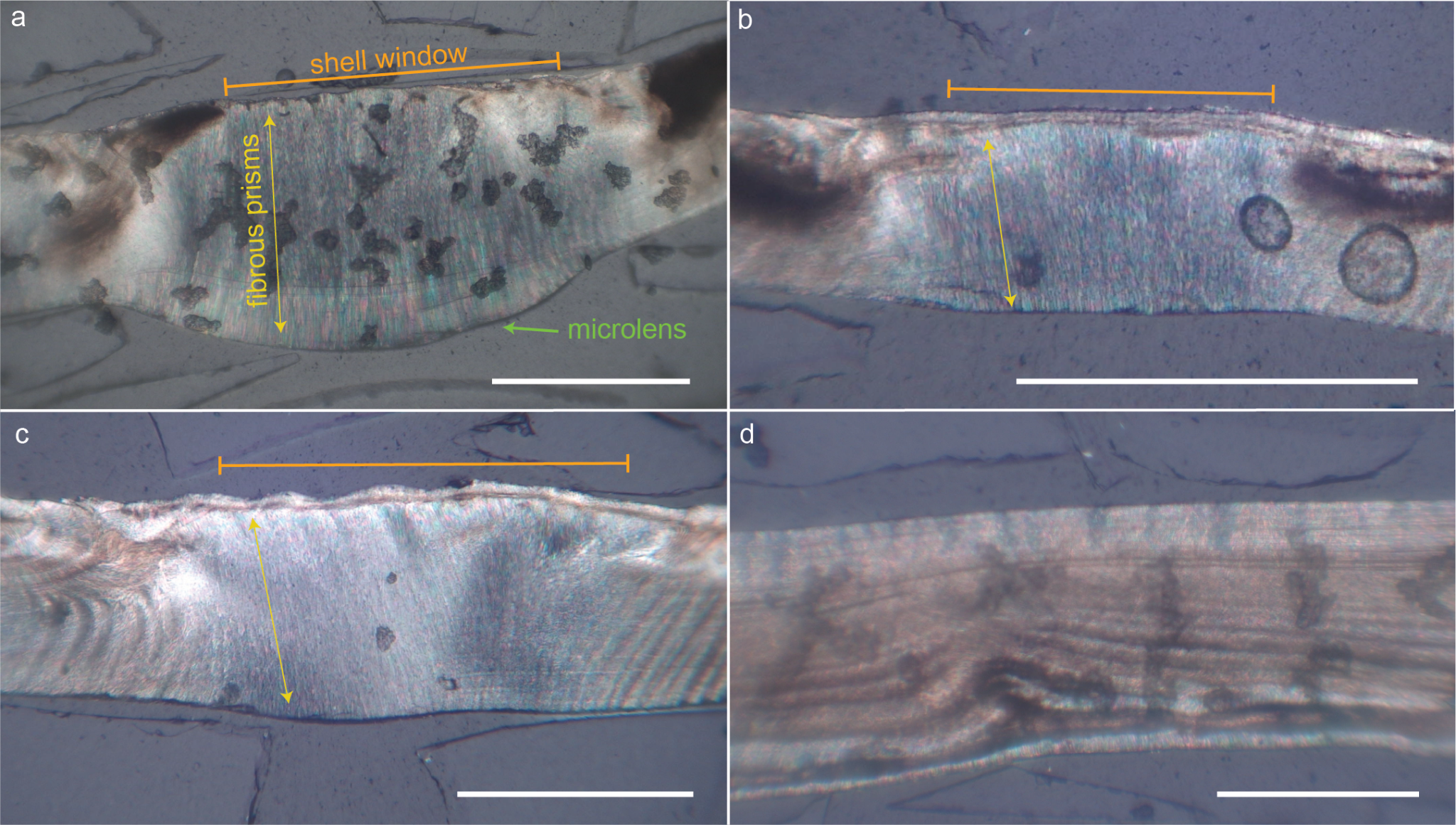

**Supplementary Figure 6: Fibrous prisms of aragonite comprise, on average, 83% of a shell’s cross-sectional thickness.** Orange bar indicates window region of shell, and yellow arrows indicate the direction of the co-oriented aragonite fiber optic bundles. These photographs were taken with a birefringent microscope, showing polished cross-sections of **(a)** shell window with microlens (green text), **(b)** polished shell window, **(c)** polished shell window, and **(d)** polished cross-section of sand-facing side of shell with no windows. Dark spots are pigmentary and other inclusions in the shells. All scale bars are 0.5 mm.

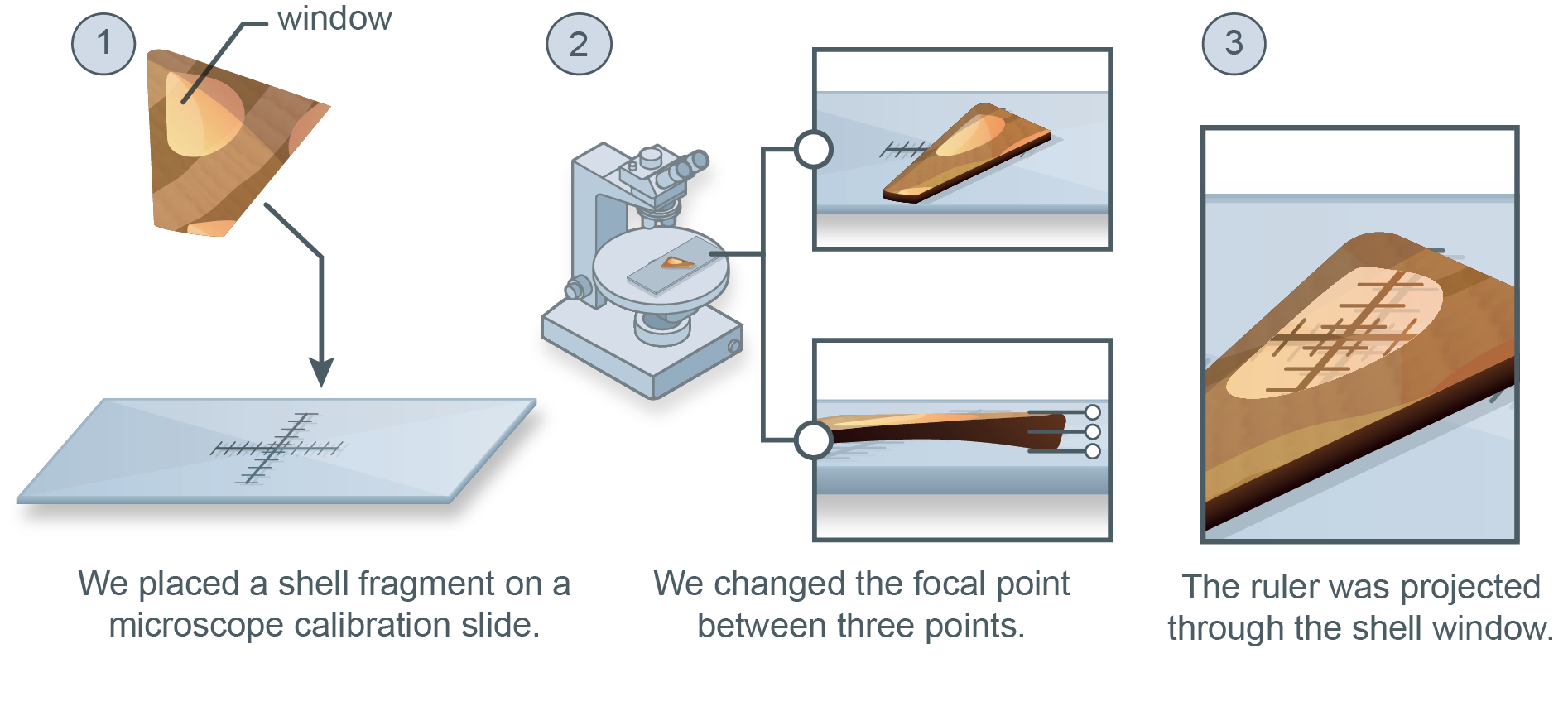

**Supplementary Figure 7. We experimentally tested image projection through a polished fragment of shell.** To experimentally test whether the windows act like fiber optic cable bundles (Figure 4), we placed a small 0.3mm thick polished fragment of shell from a heart cockle on top of a glass calibration slide. We focused the microscope on the ruler on the glass calibration slide and then adjusted the focus to refocus on the surface of the shell (~0.3mm higher than the background glass slide). Illustration is credited to Nuria Melisa Morales Garcia.

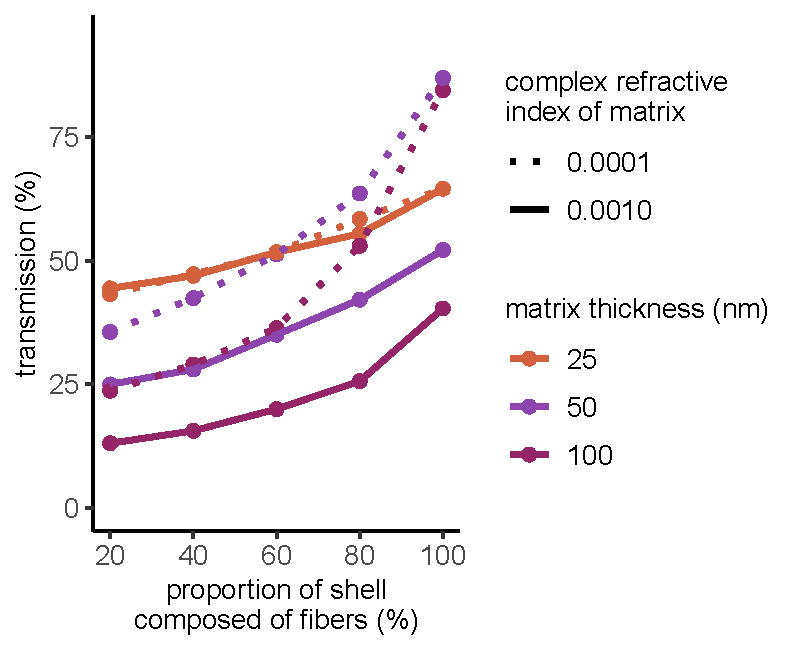

**Supplementary Figure 8: Sensitivity analyses show that results from simulations of the aragonite fiber optics are robust to variations in the refractive index and width of the organic matrix.** Here we plot the results of FDTD simulations where we varied the proportion of the shell that was composed of fibers (rather than planar aragonite), as in Figure 6a. The results are consistent across variations in the complex refractive index of the matrix and thickness of the matrix. That is, the higher the proportion of a shell composed of fibers, the greater the transmission of light through the shell. Source Data for Supplementary Figure 8 can be found in COMSOL_VaryRefractiveIndex_Simulation_Results_V11.csv (Supplementary Data 1.zip)

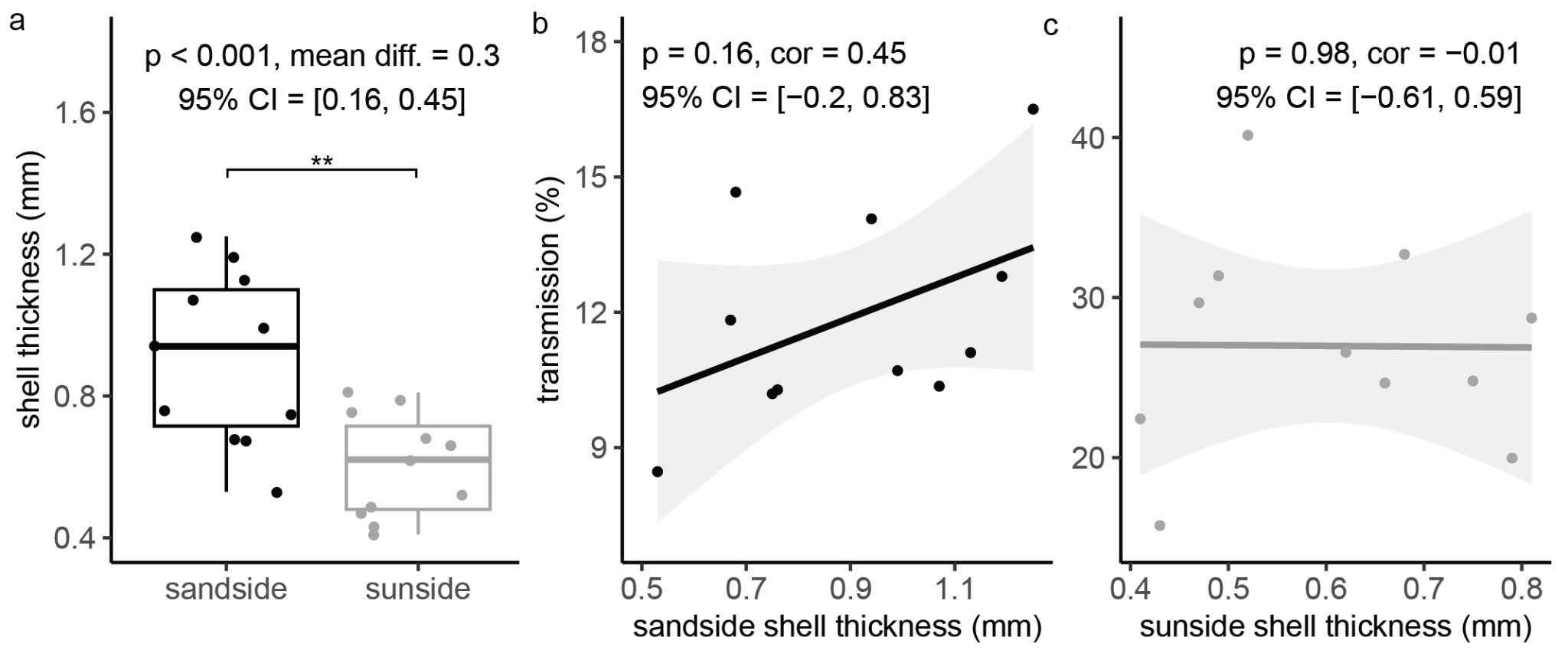

**Supplementary Figure 9. Across different individuals, shell thickness does not correlate with light transmission (300-700nm), suggesting that the optical properties of windows play an essential role in transmitting sunlight for photosynthesis.** We measured shell thickness on the sand-facing and sun-facing side of n=11 shells and tested the hypothesis that thinner-shelled individuals transmit more light. Full curves are available in Supplementary Figure 2. **(a)** The sun-facing sides of shells are significantly thinner than the sand-facing sides of shells (two-sample two-sided paired t-test, p = 0.00091, mean diff. = 0.3, 95% CI = [0.16, 0.45]). **(b)** For the sand-facing side of shells, shell thickness does not correlate significantly with mean transmission 300-700nm (Pearson's product-moment correlation test, p = 0.16, cor = 0.45, 95% CI = [-0.2, 0.83]). Indeed, the direction of the non-significant correlation is the opposite of what we expected. **(c)** For the sun-facing side of shells, shell thickness does not correlate with mean transmission 300-700 nm (Pearson’s product-moment correlation test,, p = 0.98, cor = -0.01, 95% CI = [-0.61, 0.59]. We calculated mean transmission across 300nm - 700nm. Source Data for Supplementary Figure 9 can be found in Corculum_specimens.csv (Supplementary Data 1.zip)
