## Supplementary Data 1 for "Heart cockle shells transmit sunlight to photosymbiotic algae using bundled fiber optic cables and condensing lenses": HeartCockles_AllAnalysis.R

```
## this code is all analysis for McCoy et al.  
## Heart cockle shells transmit sunlight for photosynthesis using bundled 
## fiber optic cables and condensing lenses 


### load packages
library(dplyr)
library(tidyr)
library(ggplot2)
library(stringr)
library(lsr)

# define working directory
dir <-'C:/Users/dakot/Dropbox/PC/Documents/Bivalve Optics/Analysis'

setwd(dir)

# read data files
specimen_list<-read.csv("./Corculum_specimens.csv")
data_trans<-read.csv("./UV Vis Data/UVVisTransmission_Corculumcardissa_16Mar2022.csv")
comsol_sims<-read.csv("./COMSOL/COMSOL_VaryRefractiveIndex_Simulation_Results_V11.csv")
lumerical_pvc_sims<-read.csv("./Lumerical/Lumerical_planar_vs_columnar_simulations_V2.csv")
pillarwidth_sims<-read.csv("./Lumerical/Lumerical_vary_pillarwidth_simulations_V1.csv")
FTIR_files<-list.files(path="./FTIR/FTIRMeasurements_29April2022/",pattern="*.csv",
                       ignore.case=T)
filenames_lensing<-list.files(path="./Lumerical/lensing_simulations/lensing_efield_results/",
                              pattern="*.txt")
pvc_files<-list.files(path='C:/Users/dakot/Dropbox/PC/Documents/Bivalve Optics/Analysis/Lumerical/planarvscolumnar',
                      pattern="*.txt",
                      ignore.case=T)
polishedwindows_trans<-read.csv("./Spectrophotometry_polishedwindows/Transmission_Polishedwindows_Corculum-cardissa_18Aug2023.csv")%>%
  filter(wavelength>399 & wavelength<701)
abs<-read.csv("./Absorbance Data 7May2024/Corculum-cardissa_absorbance_7May2024.csv")

head(data)
head(data_trans)

# set font size etc for my ggplots
cody_theme<- theme_classic()+
  theme(legend.position="none",
        axis.text.x=element_text(size=10),
        axis.text.y=element_text(size=10),
        axis.title.x=element_text(size=10),
        axis.title.y=element_text(size=10),
        plot.title = element_text(size=10)) 

## clean up transmission dataset
specdata_multiplemeasurements<-
  data_trans%>%
  select(-baseline_100,-baseline_0)%>%
  gather(specimen,transmission,-wavelength)%>%
  separate(specimen, c("catalog_number", "species","side",
                       "fragment","direction","measurement"), "_")
  
specdata<-data_trans%>%
  select(-baseline_100,-baseline_0)%>%
  gather(specimen,transmission,-wavelength)%>%
  separate(specimen, c("catalog_number", "species","side",
                       "fragment","direction","measurement"), "_")%>%
  group_by(catalog_number,wavelength,fragment,direction,side)%>%
  summarize(transmission=mean(transmission))%>%
  ungroup()

## save colors
orange="#d3613e"
pinkpurple="#942667"
darkpurple="#8E44AD"


A_color=orange
B_color=darkpurple
C_color=pinkpurple

###################################################################
##
##
## Plot transmission through individual polished windows
##
##
###################################################################
head(polishedwindows_trans)
paste0("Individual polished shell windows transmit ",
       round(min(polishedwindows_trans$transmission),0), "%-",
       round(max(polishedwindows_trans$transmission),0),"% of photosynthetically-active radiation (mean=",
       round(mean(polishedwindows_trans$transmission),0),"%).")

## prepare ggplot element, a shaded rectangle to how observed values
lower_observed<-mean(polishedwindows_trans$transmission)-
  sd(polishedwindows_trans$transmission)
higher_observed<-mean(polishedwindows_trans$transmission)+
  sd(polishedwindows_trans$transmission)

ggrectangle_observed<-geom_rect(aes(xmin = -Inf, xmax = Inf, ymin = lower_observed, ymax = higher_observed), fill = "gray90", alpha = 0.2)

## window 1
window1<-polishedwindows_trans%>%
  filter(specimen=="Polishedwindow1")
paste0("Window 1: ",
       round(min(window1$transmission),0), "-",
       round(max(window1$transmission),0), "% (mean=",
       round(mean(window1$transmission),0),"%).")


## window 2
window2<-polishedwindows_trans%>%
  filter(specimen=="Polishedwindow3")
paste0("Window 2: ",
       round(min(window2$transmission),0), "-",
       round(max(window2$transmission),0), "% (mean=",
       round(mean(window2$transmission),0),"%).")

ggplot(data=polishedwindows_trans)+
  # geom_point(aes(x=wavelength,y=transmission))+
  geom_smooth(aes(x=wavelength,y=transmission,color=scan),
              method="loess",span=.1,se=F)+
  scale_color_manual(values=c("black","black","black",
                              "grey70","grey70","grey70"))+
  ylab("transmission (%)")+xlab("wavelength (nm)")+
  coord_cartesian(ylim=c(20,60),xlim=c(400,700))+
  theme_classic()+
  cody_theme

ggsave(paste0("./Transmission_Graphs/","transmission_polished_windows.pdf"),
       width=3,height=3,
       units="in")

###################################################################
##
##
## Plot absorbance through cut fragments of shell
## in integrating sphere
##
##
###################################################################
head(abs)

#make long format
abs_plot<-abs%>%
  select(-Baseline.100.T)%>%
  gather(specimen,absor,-wavelength)%>%
  separate(specimen, c("catalog_number", "side","notes"), "_")%>%
  filter(wavelength>299)%>%
  filter(wavelength<700)%>%
  filter(absor<1)%>%
  # several values are indinstiguishable from 0 but negative; here
  # we convert those values to 0
  mutate(absor = if_else(absor < 0, 0, absor))
  

# plot

ggplot(data=abs_plot)+
  # add negligable amount to plot on log scale
  geom_smooth(aes(x=wavelength,y=(absor*100+.01),color=catalog_number,linetype=side),
              method="loess",span=.1,se=F)+

  ylab("absorption (%)\n in integrating sphere")+xlab("wavelength (nm)")+
  # facet_wrap("catalog_number")+
  # colorblind friendly palette
  scale_color_manual(values=c("#4477AA", "#66CCEE","#228833", "#CCBB44",
                              "#EE6677",  "#AA3377","#BBBBBB"))+
  guides(linetype = guide_legend(override.aes = list(color = "black")))+
  cody_theme+
  scale_y_continuous(trans = "log10",
                     breaks = c(0,1,5, 10,50, 100)) +
  theme(legend.position = "right")

ggsave( filename="Absorbance_logscale.pdf",
        height=3,width=4)

ggplot(data=abs_plot)+
  # add negligable amount to plot on log scale
  geom_smooth(aes(x=wavelength,y=(absor*100),color=catalog_number,linetype=side),
              method="loess",span=.1,se=F)+
  # scale_color_manual(values=c("black","black","black",
  #                             "grey70","grey70","grey70"))+
  ylab("absorption (%)\n in integrating sphere")+xlab("wavelength (nm)")+
  scale_color_manual(values=c("#4477AA", "#66CCEE","#228833", "#CCBB44",
                              "#EE6677",  "#AA3377","#BBBBBB"))+
  guides(linetype = guide_legend(override.aes = list(color = "black")))+
  # facet_wrap("catalog_number")+
  cody_theme+
  theme(legend.position = "right")

ggsave( filename="Absorbance.pdf",
        height=3,width=4)
  

# calculate absorption by UV versus PAR
abs_data<-abs_plot%>%
  mutate(absor=100*absor)

# sandside
abs_sand_UV<-abs_data%>%
  filter(side=="sandside")%>%
  filter(wavelength>299.99 & wavelength<400.01)

abs_sand_PAR<-abs_data%>%
  filter(side=="sandside")%>%
  filter(wavelength>399.99 & wavelength<700.01)

abs_sand_red<-abs_data%>%
  filter(side=="sandside")%>%
  filter(wavelength>599.99 & wavelength<700.01)

# sunside
abs_sun_UV<-abs_data%>%
  filter(side=="sunside")%>%
  filter(wavelength>299.99 & wavelength<400.01)

abs_sun_PAR<-abs_data%>%
  filter(side=="sunside")%>%
  filter(wavelength>399.99 & wavelength<700.01)

abs_sun_red<-abs_data%>%
  filter(side=="sunside")%>%
  filter(wavelength>599.99 & wavelength<700.01)

#sandside
print(paste0("The sand-facing side absorbed, on average, ",
             round(mean(abs_sand_UV$absor),0),
             "% of 300-400nm UV radiation (range = ", 
             round(min(abs_sand_UV$absor),0),
             "-",
             round(max(abs_sand_UV$absor),0),
            "%) compared to only ",
            round(mean(abs_sand_PAR$absor),0),
            "% of 400-700nm photosynthetically-active radiation (range = ", 
            round(min(abs_sand_PAR$absor),0),
            "-",
            round(max(abs_sand_PAR$absor),0),
            "%)."))

#sunside
print(paste0("The sun-facing side absorbed, on average, ",
             round(mean(abs_sun_UV$absor),0),
             "% of 300-400nm UV radiation (range = ", 
             round(min(abs_sun_UV$absor),0),
             "-",
             round(max(abs_sun_UV$absor),0),
             "%) compared to only ",
             round(mean(abs_sun_PAR$absor),0),
             "% of 400-700nm photosynthetically-active radiation (range = ", 
             round(min(abs_sun_PAR$absor),0),
             "-",
             round(max(abs_sun_PAR$absor),0),
             "%)."))

# red
print(paste0("Shells absorbed almost no 600-700nm red light, averaging ",
             round(mean(abs_sun_red$absor),0),
             "% sunside and ",
             round(mean(abs_sand_red$absor),0),
             "% sandside."))

###################################################################
##
##
## COMSOL simulations
## 
##
##
###################################################################
comsol_sims<-read.csv("./COMSOL/COMSOL_VaryRefractiveIndex_Simulation_Results_V11.csv")
head(comsol_sims)

unique(comsol_sims$matrix_spacing_nm)

# comsol_sims%>%
#   group_by(simulation_file,wavelength,matrix_spacing_nm)%>%
#   summarize(n=n())%>%
#   View()

################# transmission by orientation
sims2<-comsol_sims%>%
group_by(wavelength,Z_index,complex_refractice_index_organic_matrix)%>%
  summarize(trans_300=100*mean(transmission_300microns))

ggplot(data=sims2,
       aes(x=as.factor(Z_index),y=trans_300))+
  ggrectangle_observed+
  geom_boxplot(outlier.color="white",color=pinkpurple)+
  geom_jitter(size=0.25)+
  ylab('transmission (%)')+xlab(expression("pillar orientation (n"[z-axis]*")"))+
  theme_classic()+
  theme(axis.text.x=element_text(size=10),
        axis.text.y=element_text(size=10),
        axis.title.x=element_text(size=10),
        axis.title.y=element_text(size=10),
        plot.title = element_text(size=10),
        legend.title = element_text(size=10),
        # get rid of legend for final save
        # legend.position = "none",
        legend.text =element_text(size=10))+
  coord_cartesian(ylim=c(0,88))


ggsave( filename="Vary_Refractive_Index.pdf",
       height=2.75,width=2)

head(sims2)

wide<-sims2%>%spread(Z_index,trans_300)

head(wide)

wide$'1.53'
wide$'1.681'


tt<-t.test(wide$'1.686',wide$'1.53', paired = TRUE, alternative = "two.sided")
cohens_d<-cohensD(wide$'1.686',wide$'1.53')
print(paste0("(p=",signif(tt$p.value,2), 
             ", 95% CI=[",signif(tt$conf.int[1],3),", ",signif(tt$conf.int[2],3),"]",
             ", mean diff. = ", signif(tt$estimate,3), 
             ", degrees of freedom = ",tt$parameter[["df"]],
             ", t = ", round(tt$statistic,2),
             ", Cohen's d = ", round(cohens_d,2),
             ")"))
tt<-t.test( wide$'1.681',wide$'1.53',paired = TRUE, alternative = "two.sided")
cohens_d<-cohensD(wide$'1.681',wide$'1.53')
print(paste0("(p=",signif(tt$p.value,2), 
             ", 95% CI=[",signif(tt$conf.int[1],3),", ",signif(tt$conf.int[2],3),"]",
             ", mean diff. = ", signif(tt$estimate,3), 
             ", degrees of freedom = ",tt$parameter[["df"]],
             ", t = ", round(tt$statistic,2),
             ", Cohen's d = ", round(cohens_d,2),")"))
tt<-t.test(wide$'1.681', wide$'1.686',paired = TRUE, alternative = "two.sided")
cohens_d<-cohensD(wide$'1.681',wide$'1.686')
print(paste0("(p=",signif(tt$p.value,2), 
             ", 95% CI=[",signif(tt$conf.int[1],3),", ",signif(tt$conf.int[2],3),"]",
             ", mean diff. = ", signif(tt$estimate,3), 
             ", degrees of freedom = ",tt$parameter[["df"]],
             ", t = ", round(tt$statistic,2),
             ", Cohen's d = ", round(cohens_d,2),")"))

###################################################################
##
##
## FTIR
## 
##
##
###################################################################

## make wavenumber descending and 
## normalize curves to highest point
head(FTIR_files)

FTIR_files<-list.files(path="./FTIR/FTIRMeasurements_29April2022/",pattern="*.csv",
                       ignore.case=T)

for (i in 1:length(FTIR_files)) {

dat<-read.csv(paste0("./FTIR/FTIRMeasurements_29April2022/",FTIR_files[i],sep=""))
colnames(dat)<-c("wavenumber","R")

dat<-dat%>%filter(wavenumber>680)

## normalize the reflectance values so the maximum peak height is 100
## it is higher at times because we normalized the  FTIR measurements to
## a gold background
scale_factor<-100/max(dat$R)

dat_scaled<-dat%>%mutate(R=scale_factor*R)

##  Plot and color each region by whether that peak indicates
## A, B, or C orientation of the crystals
ggplot(data=dat_scaled)+
  ## A color (B3U)
  annotate("rect", xmin = 853, xmax = 877, ymin = -Inf, ymax = Inf,
           alpha = 0.5,fill = A_color)+
  annotate("rect", xmin = 838, xmax = 842, ymin = -Inf, ymax = Inf,
           alpha = 0.5,fill = A_color)+
  annotate("rect", xmin = 1082, xmax = 1092, ymin = -Inf, ymax = Inf,
           alpha = 0.5,fill = A_color)+
  # ## B color (B2U)
    annotate("rect", xmin = 1443, xmax = 1561, ymin = -Inf, ymax = Inf,
           alpha = 0.5,fill = B_color)+
    annotate("rect", xmin = 699, xmax = 701, ymin = -Inf, ymax = Inf,
             alpha = 0.5,fill = B_color)+
  # ## C color (B1U)
  annotate("rect", xmin = 1466, xmax = 1588, ymin = -Inf, ymax = Inf,
           alpha = 0.5,fill = C_color)+
    annotate("rect", xmin = 713, xmax = 716, ymin = -Inf, ymax = Inf,
           alpha = 0.5,fill = C_color)+
  annotate("rect", xmin = 1082, xmax = 1092, ymin = -Inf, ymax = Inf,
           alpha = 0.5,fill = C_color)+
  geom_line(aes(x=wavenumber,y=R))+
  xlab(expression ("wavenumber in "~cm^-1))+
  ylab("reflectance (normalized)")+
  # scale_x_reverse(limits = c(1800, 680))+
  scale_x_reverse()+
  cody_theme+
  theme_classic()

graph_label<-strsplit(FTIR_files[i],".CSV")[[1]][1]

### file 16 polishedshell2_window_spot3_solid.CSV
ggsave(path="./FTIR/FTIR_Figures",paste0(graph_label,".pdf",sep=""),
       width=3,height=1.5)
}


###################################################################
##
##
## Lensing simulations
## 
##
##
###################################################################

filenames_lensing<-list.files(path="./Lumerical/lensing_simulations/lensing_efield_results/",
                      pattern="*.txt")
filenames_lensing2<-  c("overallwindows-smal-inair_365nm_XZ_1_13Sep2023.txt" ,
                        "overallwindows-smal-inair_365nm_XZ_1B_13Sep2023.txt",
                        "overallwindows-smal-inair_365nm_YZ_1A_13Sep2023.txt",
                        "overallwindows-smal-inair_365nm_YZ_5_13Sep2023.txt" )
# plot a nice plot of the lensing for all the files
# for(filename in filenames_lensing) {
for(filename in filenames_lensing2) {

  

  # get the metadata from the filename
  lens<-strsplit(filename,"_")[[1]][1]
  wavelength<-strsplit(filename,"_")[[1]][2]
  axes<-strsplit(filename,"_")[[1]][3]
  
  ## get the axis names (Y and Z? X and z?). tolower() makes it lowercase
  axisA<-tolower(strsplit(axes,"")[[1]][1])
  axisB<-tolower(strsplit(axes,"")[[1]][2])
  
  # read the data
  dat<-read.csv(paste("./Lumerical/lensing_simulations/lensing_efield_results/",
                      filename,sep=""),
                sep=" ",row.names=NULL)
  head(dat)
  
  # as a sanity check, get the automatic metadata from Lumerical
  metadata_all<-paste(colnames(dat), collapse = '')
  
  # rename columns
  colnames(dat)<-c("hi","there")
  
  #find row where list of A-axis coordinates starts
  Astart<-as.numeric(rownames(dat[grepl(axisA, dat$hi), ]))+1
  Astart
  
  #find row where list of B-axis coordinates starts
  Bstart<-as.numeric(rownames(dat[grepl(axisB, dat$hi), ]))+1
  Bstart
  
  
  # get where x coordinates end
  Aend<-Bstart-2
  
  # find row where actual data starts
  Bend<-as.numeric(rownames(dat[grepl('monitor', dat$hi), ]))-1
  
  #monitor data start
  monitor_start<-Bend+5
  
  A_coords<-as.numeric(dat[Astart:Aend,"there"])
  B_coords<-as.numeric(dat[Bstart:Bend,"there"])
  
  head(A_coords)
  tail(A_coords)
  head(B_coords)
  tail(B_coords)
  
  # get spans of A and B
  Aspan<-max(A_coords)-min(A_coords)
  Bspan<-max(B_coords)-min(B_coords)
  
  ### get monitor data o the electric field
  monitor_dat<-read.csv(paste("./Lumerical/lensing_simulations/lensing_efield_results/",
                              filename,
                              sep=""),
                        skip=monitor_start,
                        sep=" ",
                        row.names = NULL,
                        header=FALSE,
                        fill=TRUE)
  head(monitor_dat)
  monitor_dat$V1<-seq(1:nrow(monitor_dat))
  
  ## now monitor dat is in the right format to be plotted!!
  
  # label the first axis's points by their actual location in microns
  dat_plotting<-monitor_dat%>%
    rename(.,A=V1)%>%
    gather(key="B",value="E_field",-1)
  
  head(dat_plotting)
  str(dat_plotting)
  
  # label the second axis's points by their actual location in microns
  dat_plotting$B<-str_replace(dat_plotting$B, str_sub(dat_plotting$B, 0, 1), '')
  head(dat_plotting)
  
  # make sure the coordinates are read as numbers
  dat_plotting$B<-as.numeric(dat_plotting$B)
  dat_plotting$A<-as.numeric(dat_plotting$A)
  
  # label the Z points numerically by their actual location in microns
  B_adjustment=Bspan/max(dat_plotting$B)
  A_adjustment=Aspan/max(dat_plotting$A)
  
  window_final<-dat_plotting%>%
    mutate(B=abs(B*B_adjustment-Bspan)*-100000000)%>%
    mutate(A=abs(A*A_adjustment-Aspan)*-100000000)
  
  min(window_final$A)
  max(window_final$A)
  
  min(window_final$B)
  max(window_final$B)
  
  # plot our window
  ggplot(window_final
           # remove top mm
           # %>%filter(B< -1000)
           ,
         aes(A/1000, B/1000, fill= E_field^2)) + 
    geom_tile() +
    scale_fill_viridis_c(option='B',
                         begin=0.1,
                         name=(expression(paste("|E|")^2))
                         # enter scale limits from infosheet
                         # ,limits=c(scale_min, scale_max)
    )+
    ## label with filename and metadata for illustrator purposes later
    labs(title=lens,subtitle = bquote(lambda == .(wavelength)))+
    # theme_classic()+
    theme_void()+
    theme(plot.title = element_text(size=6,hjust = 0.5),
          plot.subtitle = element_text(size=10,hjust = 0.5),
          legend.position="bottom",
          legend.justification = c(0.5,-1),
          legend.key.height = unit(0.15, "in"),
          legend.key.width= unit(0.15, "in"),
          legend.title.align=0.5,
          legend.text=element_text(size=10,vjust=0),
          # axis.text.x=element_text(size=10),
          # axis.text.y=element_text(size=10),
          axis.title.x=element_blank(),
          axis.title.y=element_blank())+
    coord_fixed()
  
    # coord_fixed()
  
  ggsave(paste0("./Lumerical/lensing_simulations/lensing_figures/",
                ## keep filename minus .txt
                strsplit(filename,".txt")[[1]][1],
                ".pdf"),
         width=2,height=6,
         # high resolution or not
         dpi=72,
         units="in",
         useDingbats=F)

  # ggsave(paste0("./Lumerical/lensing_simulations/lensing_figures/",
  #               ## keep filename minus .txt
  #               strsplit(filename,".txt")[[1]][1],
  #               ".png"),
  #        width=3,height=4,
  #        units="in")
}

###################################################################
##
##
## Varying pillar orientation (proportion planar vs columnar)
## Lumerical
## 
##
##
###################################################################

mypath_pvc<-'C:/Users/dakot/Dropbox/PC/Documents/Bivalve Optics/Analysis/Lumerical/planarvscolumnar'
# setwd(newdir)
pvc_files<-list.files(path=mypath_pvc,
                      pattern="*.txt",
                      ignore.case=T)

pvc_files

for (i in 1:length(pvc_files)) {
axisA="x"
axisB="y"

file_label<-strsplit(pvc_files[i],".txt")[[1]][1]
axis_description<-strsplit(file_label,"_")[[1]][4]

# if(i %in% c(11,12,13,14,15,24,30))
#   axisB="z"

if(axis_description=="transmissionlongaxis")
  axisB="z"

dat<-read.csv(paste0(mypath_pvc,"/",pvc_files[i]),
              sep=" ",row.names=NULL)
head(dat)
metadata_all<-paste(colnames(dat), collapse = '')

colnames(dat)<-c("hi","there")

#find row where list of x coordinates starts
Astart<-as.numeric(rownames(dat[grepl(axisA, dat$hi), ]))+1
Astart

#find row where list of y coordinates starts
Bstart<-as.numeric(rownames(dat[grepl(axisB, dat$hi), ]))+1
Bstart

# get where x coordinates end
Aend<-Bstart-2

# find row where actual data starts
Bend<-as.numeric(rownames(dat[grepl('transmission', dat$hi), ]))-1

#monitor data start
monitor_start<-Bend+5

A_coords<-as.numeric(dat[Astart:Aend,"there"])
B_coords<-as.numeric(dat[Bstart:Bend,"there"])

head(A_coords)
tail(A_coords)
head(B_coords)
tail(B_coords)

# get spans of A and B
Aspan<-max(A_coords)-min(A_coords)
Bspan<-max(B_coords)-min(B_coords)

### get monitor data
monitor_dat<-read.csv(paste0(mypath_pvc,"/",pvc_files[i]),
                      skip=monitor_start,
                      sep=" ",
                      row.names = NULL,
                      header=FALSE,
                      fill=TRUE)
head(monitor_dat)
monitor_dat$V1<-seq(1:nrow(monitor_dat))


## now monitor dat is in the right format to be plotted!!

# label the x-points by their actual location in microns
dat_plotting<-monitor_dat%>%
  rename(.,A=V1)%>%
  gather(key="B",value="E_field",-1)

head(dat_plotting)
str(dat_plotting)

# label the Z points numerically by their actual location in microns
dat_plotting$B<-str_replace(dat_plotting$B, str_sub(dat_plotting$B, 0, 1), '')
head(dat_plotting)

# make sure the coordinates are read as numbers
dat_plotting$B<-as.numeric(dat_plotting$B)
dat_plotting$A<-as.numeric(dat_plotting$A)

# label the Z points numerically by their actual location in microns
B_adjustment=Bspan/max(dat_plotting$B)
A_adjustment=Aspan/max(dat_plotting$A)

window_final<-dat_plotting%>%
  mutate(B=(B*B_adjustment-Bspan)*1000000)%>%
  mutate(A=abs(A*A_adjustment-Aspan)*1000000)

min(window_final$A)
max(window_final$A)

# plot our window
ggplot(window_final,aes(A, B, fill= E_field^2)) + 
  geom_tile() +
  scale_fill_viridis_c(option='B',
                       begin=0.1,
                       name=(expression(paste("|E|")^2))
                       # enter scale limits from infosheet
                       ,limits=c(0, 8)
  )+
  ## label with filename and metadata for illustrator purposes later
  labs(title=file_label)+
  theme_classic()+
  theme(axis.text.x=element_text(size=10),
        axis.text.y=element_text(size=10),
        axis.title.x=element_text(size=10),
        axis.title.y=element_text(size=10),
        plot.title = element_text(size=6),
        plot.subtitle = element_text(size=6))+
  coord_fixed()

ggsave(paste0(mypath_pvc,
              "./PVCfigures/",
              file_label,".pdf"),
       width=5,height=5,
       units="in",
       useDingbats=F)

ggsave(paste0(mypath_pvc,
              "./PVCfigures/",
              file_label,".png"),
       width=5,height=5,
       units="in")

}


###################################################################
##
## now simple graph of proportion transmission varying
##
###################################################################

head(lumerical_pvc_sims)

pvc<-lumerical_pvc_sims%>%
  select(matrix_width_group,
        proportion_planar,
        complex_refractive_index_K,
        transmission_300microns)%>%
  group_by(matrix_width_group,proportion_planar,complex_refractive_index_K)%>%
  mutate(transmission=mean(transmission_300microns))

pvc$complex_refractive_index_K<-as.factor(format(pvc$complex_refractive_index_K,scientific = FALSE))
pvc$matrix_width_group<-as.factor(pvc$matrix_width_group)

head(pvc)

## just one micron for main text
ggplot(pvc%>%
         filter(matrix_width_group=="100",
                complex_refractive_index_K=="0.0010"))+
  ggrectangle_observed+
  geom_point(aes(x=100-proportion_planar*100,y=transmission*100),size=2,
             color=pinkpurple)+
  geom_line(aes(x=100-proportion_planar*100,y=transmission*100),size=1,
            color=pinkpurple)+
  xlab("proportion of shell\ncomposed of fibers (%)")+
  ylab("transmission (%)")+
  coord_cartesian(ylim=c(12,44))+
  theme_classic()+
  theme(axis.text.x=element_text(size=10),
        axis.text.y=element_text(size=10),
        axis.title.x=element_text(size=10),
        axis.title.y=element_text(size=10),
        plot.title = element_text(size=10))+
  coord_cartesian(ylim = c(10,65))

ggsave("varyplanarvscolumnar_proportionplot.pdf",
       width=2.75,height=2.75,
       units="in",
       useDingbats=F)

## all sweeps
ggplot(pvc)+
  geom_point(aes(x=100-proportion_planar*100,y=transmission*100,
                 color=matrix_width_group),size=2)+
  geom_line(aes(x=100-proportion_planar*100,y=transmission*100,
                color=matrix_width_group,linetype=complex_refractive_index_K),size=1)+
  xlab("proportion of shell\ncomposed of fibers (%)")+
  ylab("transmission (%)")+
  scale_color_manual(values=c(orange, darkpurple,pinkpurple),
                     name = "matrix thickness (nm)")+
  scale_linetype_manual(values=c(3,1),name = "complex refractive\nindex of matrix")+
  theme_classic()+
  coord_cartesian(ylim=c(0,95))+
  theme(axis.text.x=element_text(size=10),
        axis.text.y=element_text(size=10),
        axis.title.x=element_text(size=10),
        axis.title.y=element_text(size=10),
        plot.title = element_text(size=10),
        legend.text = element_text(size=10),
        legend.title = element_text(size=10))

ggsave("varyplanarvscolumnar_proportionplot_supplement.pdf",
       width=4,height=3.25,
       units="in",
       useDingbats=F)


################################################################
################################################################
######
###### Vary pillar size widely
######
################################################################
################################################################

pw<-pillarwidth_sims%>%
  select(-random1,-random2,-random3,
         -round,-transmission_50microns,
         -transmission_300microns,-transmission_sd,
         -length_m,-width_m,-matrix_thicknes_nm)%>%
  gather(.,key="wavelength",value="transmission",-pillar_width_microns ,
         -matrix_thickness_nm,-complex_refractive_index)

head(pw)

pw$wavelength<-as.numeric(str_replace(pw$wavelength, 'wavelength_', ''))

pw<-pw%>%mutate_all(., function(x) as.numeric((x)))%>%
  group_by(pillar_width_microns)%>%
  summarize(trans=mean(transmission))

pw

ggplot(pw)+
  ggrectangle_observed+
  geom_point(aes(x=pillar_width_microns,y=trans^6*100),size=2,
             color=pinkpurple)+
  geom_line(aes(x=pillar_width_microns,y=trans^6*100),size=1,
            color=pinkpurple)+
  theme_classic()+
  xlab(expression("pillar width ("*mu*"m)"))+
  ylab("transmission (%)")+
  theme(axis.text.x=element_text(size=10),
        axis.text.y=element_text(size=10),
        axis.title.x=element_text(size=10),
        axis.title.y=element_text(size=10),
        plot.title = element_text(size=10))+
  coord_cartesian(ylim=c(0,88))
   # scale_x_continuous(trans='log10') 


ggsave("varywidthwidely.pdf",
       width=2.75,height=2.75,
       units="in",
       useDingbats=F)

###################################################################
##
##
## Plot transmission for all
##
##
###################################################################

# make list of all specimens
specimens<-unique(specdata$catalog_number)

# plot transmission for each
for (i in 1:length(specimens)) {
ggplot(data=specdata_multiplemeasurements%>%
         filter(catalog_number==specimens[i]))+
  geom_smooth(aes(x=wavelength,y=transmission,color=side,fill=side),
              method="loess",span=0.1)+
  scale_color_manual(values=c("black","grey70"))+
  scale_fill_manual(values=c("grey50","grey80"))+
  ylab("transmission (%)")+xlab("wavelength (nm)")+
  geom_text(x=350,y=60,label=paste(specimens[i]),size=2)+
  coord_cartesian(ylim=c(0,60))+
  theme_classic()+
  theme(legend.position="none",
        axis.text.x=element_text(size=10),
        axis.text.y=element_text(size=10),
        axis.title.x=element_text(size=10),
        axis.title.y=element_text(size=10),
        strip.text = element_blank(),
        strip.background = element_blank()) 
  
  ggsave(paste0("./Transmission_Graphs/",specimens[i],"_Transmission.pdf"),
         width=2,height=2,
         units="in")
}

# plot transmission for each
# for (i in 1:length(specimens)) {
#   ggplot(data=specdata%>%
#            filter(catalog_number==specimens[i]))+
#     geom_smooth(aes(x=wavelength,y=transmission,color=side),
#                 method="loess",span=0.1)+
#     scale_color_manual(values=c("black","grey70"))+
#     ylab("transmission (%)")+xlab("wavelength (nm)")+
#     geom_text(x=350,y=60,label=paste(specimens[i]),size=2)+
#     coord_cartesian(ylim=c(0,60))+
#     theme_classic()+
#     theme(legend.position="none",
#           axis.text.x=element_text(size=10),
#           axis.text.y=element_text(size=10),
#           axis.title.x=element_text(size=10),
#           axis.title.y=element_text(size=10),
#           strip.text = element_blank(),
#           strip.background = element_blank()) 
#   
#   ggsave(paste0("./Transmission_Graphs/",specimens[i],"_Transmission_noerrorbars.pdf"),
#          width=2,height=2,
#          units="in")
# }

## get range and mean transmission across all specimens

### sun-facing side, PAR
specdata_print<-specdata%>%
  filter(side=="sunside")%>%
  filter(wavelength>399.99 & wavelength <700.01)
print(paste0("Across all specimens, the sun-facing side transmits ",
             round(min(specdata_print$transmission),0),
             "-",
             round(max(specdata_print$transmission),0),
             "% of 400-700nm photosynthetically active radiation (mean = ",
             round(mean(specdata_print$transmission),0),
             "%)"))

### sun-facing side, UV
specdata_print<-specdata%>%
  filter(side=="sunside")%>%
  filter(wavelength>299.99 & wavelength <400.01)
print(paste0("but only transmits ",
             round(min(specdata_print$transmission),0),
             "-",
             round(max(specdata_print$transmission),0),
             "% of 300-400nm UV radiation (mean = ",
             round(mean(specdata_print$transmission,0)),
             "%)."))

### sand-facing side, PAR
specdata_print<-specdata%>%
  filter(side=="sandside")%>%
  filter(wavelength>399.99 & wavelength <700.01)
print(paste0("Correspondingly the sand-facing side transmits ",
             round(min(specdata_print$transmission),0),
             "-",
             round(max(specdata_print$transmission),0),
             "% of photosynthetically active radiation (mean = ",
             round(mean(specdata_print$transmission),0),
             "%)"))

### sun-facing side, UV
specdata_print<-specdata%>%
  filter(side=="sandside")%>%
  filter(wavelength>299.99 & wavelength <400.01)
print(paste0("and ",
             round(min(specdata_print$transmission),0),
             "-",
             round(max(specdata_print$transmission),0),
             "% of UV radiation (mean = ",
             round(mean(specdata_print$transmission,0)),
             "%)."))

### sun-facing side, red
specdata_print<-specdata%>%
  filter(side=="sunside")%>%
  filter(wavelength>599.99 & wavelength <700.01)
print(paste0("The sun-facing side transmits ",
             round(min(specdata_print$transmission),0),
             "-",
             round(max(specdata_print$transmission),0),
             "% of 600-700nm red light (mean = ",
             round(mean(specdata_print$transmission,0)),
             "%)."))


### get average transmission for each specimen
# for total light, UV, PAR, and red light

# make list of all specimens
specimens<-unique(specdata$catalog_number)

results<-matrix(ncol=9, nrow=length(specimens))
colnames(results)<-c("specimen",
                     "TotalTransmission_sunside",
                     "Transmission300-400_sunside",
                     "Transmission600-700_sunside",
                     "Transmission400-700_sunside",
                     "TotalTransmission_sandside",
                     "Transmission300-400_sandside",
                     "Transmission600-700_sandside",
                     "Transmission400-700_sandside"
                     )
for (i in 1:length(specimens)) {
  results[i,"specimen"]<-specimens[i]
  ## sunside
  data_sunside <- specdata%>%
    filter(catalog_number==specimens[i])%>%
    filter(side=="sunside")%>%
    select(wavelength,transmission)
  data_sunside_all<-data_sunside%>%
    filter(wavelength>299.99 & wavelength < 700.01)
  data_sunside_UV<-data_sunside%>%
    filter(wavelength>299.99 & wavelength < 400.01)
  data_sunside_PAR<-data_sunside%>%
    filter(wavelength>399.99 & wavelength < 700.01)
  data_sunside_red<-data_sunside%>%
    filter(wavelength>599.99 & wavelength < 700.01)
  results[i,"TotalTransmission_sunside"]<-mean(data_sunside_all$transmission)
  results[i,"Transmission300-400_sunside"]<-mean(data_sunside_UV$transmission)
  results[i,"Transmission600-700_sunside"]<-mean(data_sunside_red$transmission)
  results[i,"Transmission400-700_sunside"]<-mean(data_sunside_PAR$transmission)
  ## sandside
  data_sandside <- specdata%>%
    filter(catalog_number==specimens[i])%>%
    filter(side=="sandside")%>%
    select(wavelength,transmission)
  data_sandside_all<-data_sandside%>%
    filter(wavelength>299.99 & wavelength < 700.01)
  data_sandside_UV<-data_sandside%>%
    filter(wavelength>299.99 & wavelength < 400.01)
  data_sandside_PAR<-data_sandside%>%
    filter(wavelength>399.99 & wavelength < 700.01)
  data_sandside_red<-data_sandside%>%
    filter(wavelength>599.99 & wavelength < 700.01)
  results[i,"TotalTransmission_sandside"]<-mean(data_sandside_all$transmission)
  results[i,"Transmission300-400_sandside"]<-mean(data_sandside_UV$transmission)
  results[i,"Transmission600-700_sandside"]<-mean(data_sandside_red$transmission)
  results[i,"Transmission400-700_sandside"]<-mean(data_sandside_PAR$transmission)
}
results<-as.data.frame(results)

results[,2:9] <- sapply( results[,2:9], as.numeric )

# compare UV to red
results$redminusblueUV_sunside<-results$"Transmission600-700_sunside"-
  results$"Transmission300-400_sunside"

# compare UV to red
results$redminusblueUV_sandside<-results$"Transmission600-700_sandside"-
  results$"Transmission300-400_sandside"

# check how much more red than UV light is transmitted
results$reddividedbyUV_sunside<-results$"Transmission600-700_sunside"/
  results$"Transmission300-400_sunside"

results$reddividedbyUV_sandside<-results$"Transmission600-700_sandside"/
  results$"Transmission300-400_sandside"

# print some results
print(paste0("mean ratio = ",
             round(mean(results$reddividedbyUV_sunside),1),
             ", range = (",
             round(min((results$reddividedbyUV_sunside)),1),
             "-",
             round(max(results$reddividedbyUV_sunside),1),
             ")."))

#########################
## plot this stuff
#########################

head(results)

results_long<-results%>%
  gather(label,transmission,-specimen)

ggplot(data=results_long%>%
         filter(label %in% c("TotalTransmission_sunside",
                             "TotalTransmission_sandside")),
       aes(x=label,y=transmission,color=label))+
  scale_color_manual(values=c("black","gray60"))+
  xlab("shell side")+
  ylab("mean transmission (%)")+
  geom_boxplot(outlier.shape=NA, width=0.5)+
  geom_jitter(size=1)+
  coord_cartesian(ylim=c(0,41))+
  cody_theme+
  theme(axis.title.x = element_blank())

ggsave("./Transmission_Graphs/Boxplot_Sunside-vs-Sandside.pdf",
       width=3.1,height=1.8,
       units="in")

ggplot(data=results_long%>%
         filter(label %in% c("Transmission300-400_sunside",
                             "Transmission600-700_sunside")),
       aes(x=label,y=transmission,color=label))+
  scale_color_manual(values=c("purple4","red4"))+
  xlab("shell side")+
  ylab("mean transmission (%)")+
  geom_boxplot(outlier.shape=NA)+
  geom_jitter(size=1)+
  coord_cartesian(ylim=c(0,60))+
  cody_theme+
  theme(axis.title.x = element_blank())+
  theme(
        # axis.title.y=element_text(colour="grey70"),
        # axis.text.y=element_text(colour="grey70"),
        axis.ticks = element_line(colour="grey60"),
        axis.line= element_line(colour="white"))
ggsave("./Transmission_Graphs/Boxplot_Sunside_Long-vs-Short-Wavelength.pdf",
       width=1.8,height=1.8,
       units="in")

ggplot(data=results_long%>%
         filter(label %in% c("Transmission300-400_sandside",
                             "Transmission600-700_sandside")),
       aes(x=label,y=transmission,color=label))+
  scale_color_manual(values=c("purple4","red4"))+
  xlab("shell side")+
  ylab("mean transmission (%)")+
  geom_boxplot(outlier.shape=NA)+
  geom_jitter(size=1)+
  coord_cartesian(ylim=c(0,60))+
  cody_theme+
  theme(axis.title.x = element_blank())+
  theme(
    # axis.title.y=element_text(colour="grey70"),
    # axis.text.y=element_text(colour="grey70"),
    axis.ticks = element_line(colour="black"),
    axis.line= element_line(colour="white"))
ggsave("./Transmission_Graphs/Boxplot_Sandside_Long-vs-Short-Wavelength.pdf",
       width=1.8,height=1.8,
       units="in")


#######################
###paired two sample t-test
#######################
tt<-t.test(results$"TotalTransmission_sunside",
           results$"TotalTransmission_sandside",
           paired = TRUE, alternative = "two.sided")
cohens_d<-cohensD(results$"TotalTransmission_sunside",
                  results$"TotalTransmission_sandside")

print(paste0("(p=",signif(tt$p.value,2), 
             ", 95% CI=[",signif(tt$conf.int[1],3),", ",signif(tt$conf.int[2],3),"]",
             ", mean diff. = ", signif(tt$estimate,3), 
             ", degrees of freedom = ",tt$parameter[["df"]],
             ", t = ", round(tt$statistic,2),
             ", Cohen's d = ", round(cohens_d,2),")"))


tt<-t.test(results$"Transmission600-700_sunside",
           results$"Transmission300-400_sunside",
           paired = TRUE, alternative = "two.sided")
cohens_d<-cohensD(results$"Transmission600-700_sunside",
                  results$"Transmission300-400_sunside")

print(paste0("(p=",signif(tt$p.value,2), 
             ", 95% CI=[",signif(tt$conf.int[1],3),", ",signif(tt$conf.int[2],3),"]",
             ", mean diff. = ", signif(tt$estimate,3), 
             ", degrees of freedom = ",tt$parameter[["df"]],
             ", t = ", round(tt$statistic,2),
             ", Cohen's d = ", round(cohens_d,2),")"))

tt<-t.test(results$"Transmission600-700_sandside",
           results$"Transmission300-400_sandside",
           paired = TRUE, alternative = "two.sided")
cohens_d<-cohensD(results$"Transmission600-700_sandside",
                  results$"Transmission300-400_sandside")

print(paste0("(p=",signif(tt$p.value,2), 
             ", 95% CI=[",signif(tt$conf.int[1],3),", ",signif(tt$conf.int[2],3),"]",
             ", mean diff. = ", signif(tt$estimate,3), 
             ", degrees of freedom = ",tt$parameter[["df"]],
             ", t = ", round(tt$statistic,2),
             ", Cohen's d = ", round(cohens_d,2),")"))

tt<-t.test(results$redminusblueUV_sunside,
           results$redminusblueUV_sandside,
           paired = TRUE, alternative = "two.sided")
cohens_d<-cohensD(results$redminusblueUV_sunside,
                  results$redminusblueUV_sandside)


print(paste0("(p=",signif(tt$p.value,2), 
             ", 95% CI=[",signif(tt$conf.int[1],3),", ",signif(tt$conf.int[2],3),"]",
             ", mean diff. = ", signif(tt$estimate,3), 
             ", degrees of freedom = ",tt$parameter[["df"]],
             ", t = ", round(tt$statistic,2),
             ", Cohen's d = ", round(cohens_d,2),")"))


###################################################################
##
##
## Assess shell width
##
##
###################################################################
head(specdata)
head(specimen_list)

# specdata%>%
#   group_by(catalog_number)%>%
#   summarize(trans=mean(transmission))

trans<-results%>%
  select(specimen,TotalTransmission_sunside,TotalTransmission_sandside)

colnames(trans)<-c("catalog_number","trans_sunside","trans_sandside")

allspecs<-merge(trans,specimen_list,by="catalog_number")

# run t test on the hypothesis that sandside shell is different thickness than sunside shell
t_thick<-t.test(allspecs$sandside_thickness_mm , allspecs$sunside_thickness_mm, 
           paired = TRUE, 
           alternative = "two.sided")

# write pretty text
ifelse (t_thick$p.value < 0.001,
        #yes
        t_thick_text<-paste0("p < 0.001", 
             ", mean diff. = ", signif(t_thick$estimate,2),
             "\n95% CI = [",signif(t_thick$conf.int[1],2),", ",
             signif(t_thick$conf.int[2],2),"]"),
        #no
          t_thick_text<-paste0("p = ",signif(t_thick$p.value,2), 
                       ", mean diff. = ", signif(t_thick$estimate,2),
                       "\n95% CI = [",signif(t_thick$conf.int[1],2),", ",
                       signif(t_thick$conf.int[2],2),"]")
)
  

# run correlation tests on the hypothesis that the thinner the shell, the greater the transmission
cortest_sand<-cor.test(x=allspecs$sandside_thickness_mm,
                       y=allspecs$trans_sandside,
                       alternative = "two.sided")

cortest_sun<-cor.test(x=allspecs$sunside_thickness_mm,
                      y=allspecs$trans_sunside,
                      alternative = "two.sided")

# prepare nice-looking summary text
sand_cor_text<-paste0("p = ", round(cortest_sand$p.value,2),
                      ", cor = ",round(cortest_sand$estimate,2),"\n95% CI = ",
                      "[",round(cortest_sand$conf.int[1],2), ", ",round(cortest_sand$conf.int[2],2),"]",sep="")

sand_cor_text

sun_cor_text<-paste0("p = ", round(cortest_sun$p.value,2),
                     ", cor = ",round(cortest_sun$estimate,2),"\n95% CI = ",
                     #confint
                     "[",round(cortest_sun$conf.int[1],2), ", ",round(cortest_sun$conf.int[2],2),"]",sep="")

sun_cor_text

## plot and save
ggplot(data=allspecs,aes(x=sunside_thickness_mm,y=trans_sunside))+
    geom_smooth(method="lm",color="grey60",fill="grey85")+
  geom_point(color="grey65",size=1)+ 
  xlab("sunside shell thickness (mm)")+
  ylab("transmission (%)")+

  coord_cartesian(ylim=c(15,46))+
  annotate("text", 
           x = 0.8, y = 46, 
           hjust=1,
           vjust=1,
           label = sun_cor_text, 
           color = "black",
           size = 10/2.845276) +
  cody_theme

ggsave("sunside_thickness.pdf",
       width=2.5,height=3)

ggplot(data=allspecs,aes(x=sandside_thickness_mm,y=trans_sandside))+
    geom_smooth(method="lm",color="black",fill="grey85")+
  geom_point(color="black",size=1)+  
  xlab("sandside shell thickness (mm)")+
  ylab("transmission (%)")+

  coord_cartesian(ylim=c(7,18))+
  annotate("text", 
           x = 0.5, y = 18,
           hjust=0,
           vjust=1,
           label = sand_cor_text, 
           color = "black",
           size = 10/2.845276) +
  # coord_cartesian(ylim=c(6,18)) +
  cody_theme

ggsave("sandside_thickness.pdf",
       width=2.5,height=3)

ggplot(data=allspecs%>%select(catalog_number,sunside_thickness_mm,sandside_thickness_mm)%>%
         gather(., measurement, thickness, -catalog_number))+
    geom_boxplot(aes(y=thickness,x=measurement,color=measurement))+
  geom_jitter(aes(y=thickness,x=measurement,color=measurement),
              size=1)+
  scale_color_manual(values=c("black","grey65"))+
  scale_x_discrete(labels = c("sandside","sunside"))+
  xlab("")+
  ylab("shell thickness (mm)")+
  coord_cartesian(ylim=c(0.4,1.8))+
  annotate("text", 
           x=1.5,y =1.7, 
           label = t_thick_text, 
           color = "black",
           size = 10/2.845276) +
  cody_theme

ggsave("thickness.pdf",
       width=2.5,height=3)
```
